# Adaptations to water stress and pastoralism in the Turkana of northwest Kenya

**DOI:** 10.1101/2023.01.17.524066

**Authors:** AJ Lea, IV Caldas, KM Garske, J Echwa, M Gurven, C Handley, J Kahumbu, Kamau, P Kinyua, F Lotukoi, A Lopurudoi, S Lowasa, R Mallarino, D Martins, PW Messer, C Miano, B Muhoya, J Peng, T Phung, JD Rabinowitz, A Roichman, R Siford, A Stone, AM Taravella Oill, S Mathew, MA Wilson, JF Ayroles

**Affiliations:** Department of Biological Sciences, Vanderbilt University, Nashville, TN, USA; Turkana Health and Genomics Project, Mpala Research Centre, Nanyuki, Kenya; Department of Computational Biology, Cornell University, Ithaca, NY, USA; Department of Ecology and Evolutionary Biology, Princeton University, Princeton 08544, New Jersey, USA; Lewis Sigler Institute for Integrative Genomics, Princeton University, Princeton 08544, New Jersey, USA; Department of Anthropology, University of California Santa Barbara, Santa Barbara, CA, USA; School of Human Evolution and Social Change, Arizona State University, Tempe, AZ, USA; Institute of Primate Research, National Museums of Kenya, Nairobi 00502, Kenya; Department of Biochemistry, School of Medicine, University of Nairobi, Nairobi, Kenya; Department of Molecular Biology, Princeton University, Princeton 08544, New Jersey, USA; Turkana Basin Institute, Turkana, Kenya; Department of Chemistry, Princeton University, Princeton 08544, New Jersey, USA; Institute of Human Origins, Arizona State University, Tempe, AZ, USA; Center for Evolution and Medicine, Arizona State University, Tempe, AZ, USA; School of Life Sciences, Arizona State University, Tempe, AZ, USA

**Keywords:** Turkana, arid living, STC1, human evolution, pastoralism, soft-sweep, adaptation

## Abstract

The Turkana people inhabit arid regions of east Africa—where temperatures are high and water is scarce—and they practice subsistence pastoralism, such that their diet is primarily composed of animal products. Working with Turkana communities, we sequenced 367 genomes and identified 8 regions putatively involved in adaptation to water stress and pastoralism. One of these regions includes a putative enhancer for STC1—a kidney-expressed gene involved in the response to dehydration and the metabolism of purine-rich foods such as red meat. We show that STC1 is induced by antidiuretic hormone in humans, is associated with urea levels in the Turkana themselves, and is under strong selection in this population (s∼0.041). This work highlights that partnerships with subsistence-level groups can lead to new models of human physiology with biomedical relevance.

## INTRODUCTION

Humans inhabit an astounding variety of challenging habitats—from high altitude plateaus to arctic tundras to scorching deserts. In many cases, natural selection has played a key role in sustaining human populations in extreme environments (1, 2). For example, the hypoxic conditions experienced by Tibetans living at high altitudes have selected for changes in red blood cell production regulated by EPAS1 (3), while the carnivorous diets of Greenlandic Inuits have selected for changes in fatty acid metabolism regulated by the FADS gene cluster (4). Such examples highlight that, in addition to helping us understand our evolutionary history, studies of natural selection can uncover new genotype-phenotype links of biomedical importance. However, despite great interest in uncovering the genetic basis of adaptation, few studies have robustly linked ecological selection pressures, genetic variation, and human phenotypes (2).

To do so, we partnered with the Turkana people of northwest Kenya, who manage several dietary and climatic challenges (Figure 1A-B). The Turkana are an Eastern Nilotic group, and like other members of their lineage, they originated in the Nile Valley desert and began nomadic pastoralist practices ∼5-8 thousand years ago (5, 6). Oral histories suggest they migrated to Kenya in the last few hundred years, where they continue to inhabit similar climates and engage in similar lifestyles (7).

**Figure 1.**
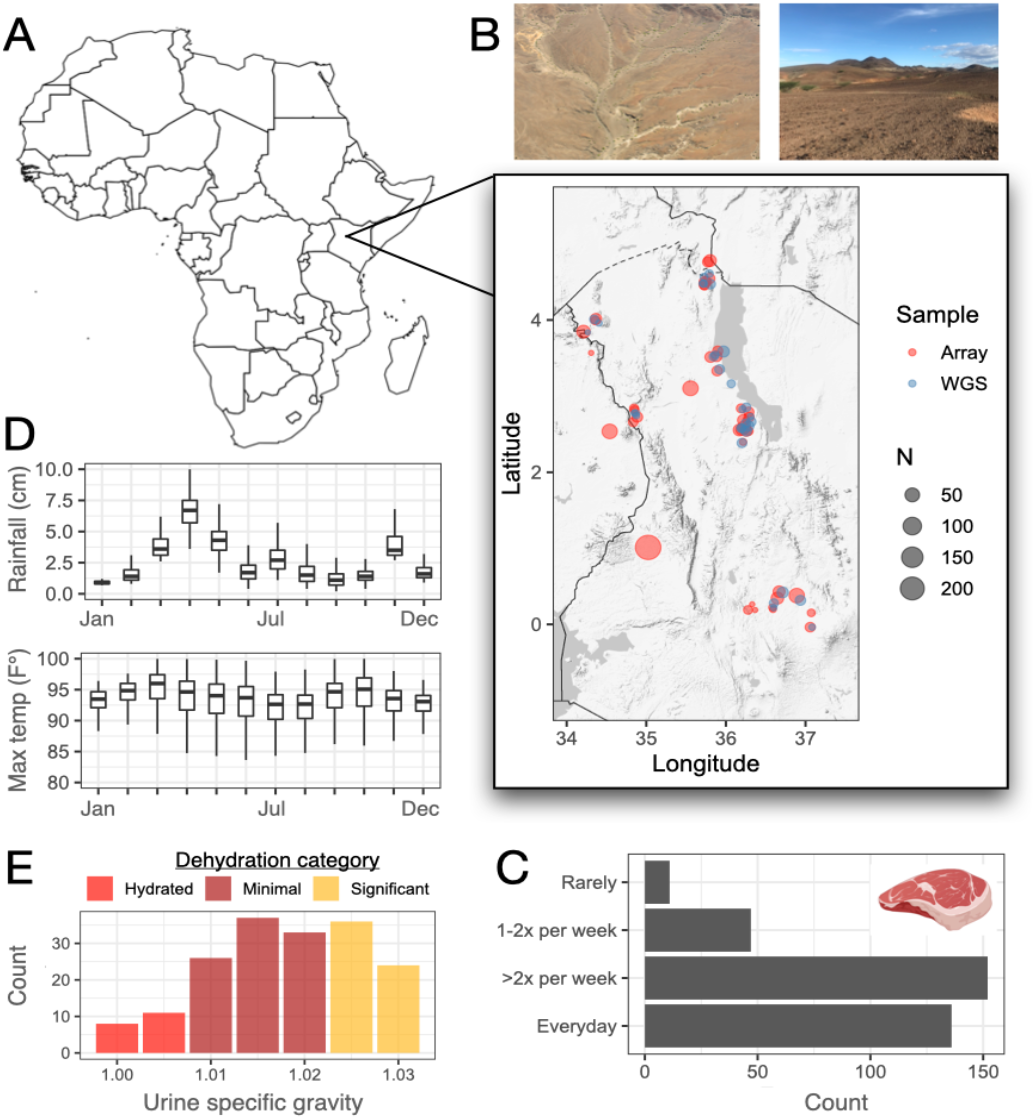
Ecology and lifestyle of the Turkana people. A) Map of Africa with an inset showing northwest Kenya (the present day homelands of the Turkana people). Dots indicate where samples were collected for this study, for both whole genome sequencing (WGS) and array genotyping. B) Representative photographs of the arid ecology of the Turkana region (photos taken by the authors). C) Number of Turkana pastoralists who consume meat at different self-reported frequencies. D) Average rainfall and maximum temperature for the Turkana country region by month. Data were sourced from WorldClim (10). E) Number of Turkana pastoralists with different urine specific gravity values, colored by whether each value meets the criteria for dehydration.

As a result of their pastoralist practices, the Turkana diet is protein-rich, with previous studies estimating that 70-80% of the diet is animal-derived (e.g., from milk, blood, marrow, and red meat) (8). In agreement, our dietary interviews revealed that 74% and 96.8% of Turkana pastoralists consumed blood and red meat several times per week (n=346; Figure 1C, Table S1). The estimated protein intake from this diet exceeds the FAO/WHO requirements by >300% (9) and would be considered atherogenic by most standards.

The Turkana practice nomadic pastoralism because they inhabit extremely arid, water-limited (Figure 1D) landscapes that cannot easily support agriculture, hunting and gathering, or other common subsistence strategies; instead, they rely on opportunistic exploitation of transient rainfall and vegetation by their livestock. To quantify the impact of water limitation, we interviewed Turkana pastoralists and found that water stress was a daily issue: 76.2% of Turkana spend more than a few hours a day collecting water, and 99.2% perceive this water to be insufficient in amount (n=311; Table S1). Using measures of urine-specific gravity, we also found that 89.1% of people meet the physiological criteria for dehydration (n=175; Figure 1E, Table S1), emphasizing the challenge of maintaining fluid balance in this environment.

### Selective sweeps in the Turkana

We hypothesized that exposure to an animal product-rich diet and an arid ecology would select for genetic variants regulating metabolism and dehydration stress in the Turkana. To test this, we scanned for signatures of selection in 308 Turkana genomes sequenced at high (>20x, n=106) and medium (∼6x, n=202) coverage. To understand population genetic parameters that could impact our analyses and interpretation, we also sequenced 59 genomes from nearby Kenyan/Ugandan groups, namely the El Molo, Ik, Karamojong, Masaai, Ngitepes (also known as Tepeth), Pokot, Rendille, and Samburu (Figure S1, Table S2-S3).

We genotyped 7,767,165 variants with a minor allele frequency >1% in our high-coverage Turkana dataset and then imputed missing data in the medium-coverage samples (Figure 2A, Figure S2). We performed extensive QC on our combined, imputed WGS dataset (Figure S3-6), including corroborating our WGS-derived genotype calls using independent calls from the Infinium Global Screening Array (R2 between WGS- and array-derived genotypes for 108 paired samples=0.96 ± 0.03; Table S4). Overall, population genetic analyses of the WGS dataset highlighted two key takeaways that were confirmed with the array dataset (n=783 individuals total) and are consistent with the literature (6, 11). First, there has been little European admixture in the East African groups we worked with (mean European ancestry proportion estimated by RFMix=6.6 ± 4.4%; Figure S7-9, Table S5). Second, though these groups are culturally distinct, most are not statistically distinguishable at the genomic level using common summary statistics (Figure 2A, Figure S5, Figure S8, Figure S10).

**Figure 2.**
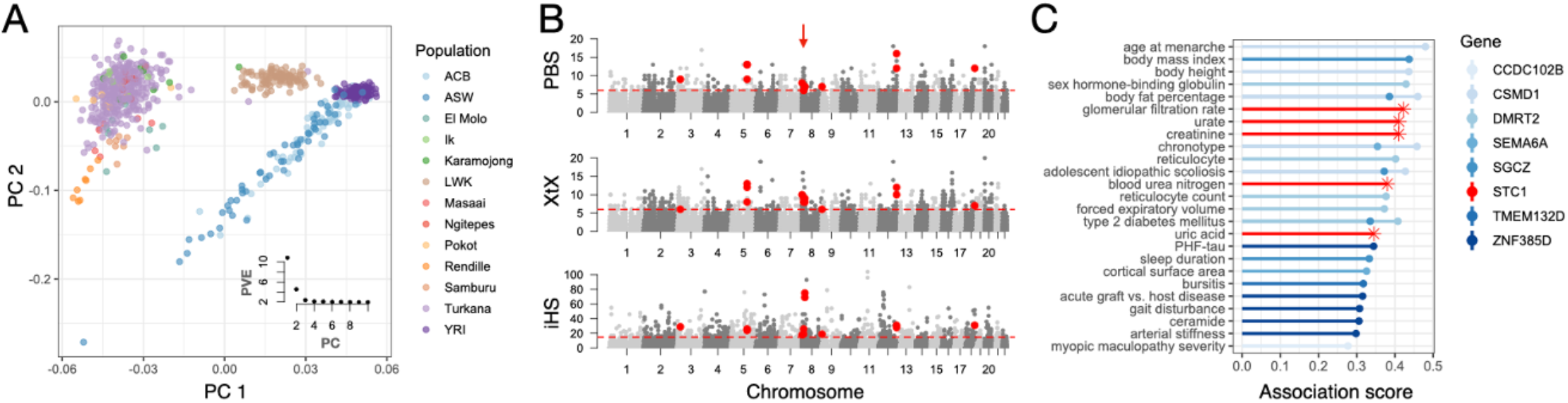
Population and evolutionary genetics. A) Principal components analysis comparing the Turkana as well as other study communities to African populations included in the 1000 Genomes Study (14). Inset shows the percent variation (PVE) explained by each principal component (PC). B) Number of PBS, XtX, and iHS outliers per 50 kb window. Red dotted lines represent the cutoff for the 99th percentile of each empirical distribution, and windows that passed this criterion for all three statistics are highlighted in red. The candidate region that falls near the STC1 gene is highlighted with a red arrow. C) Phenotypes associated with genes that fall in or near candidate regions. Phenotypic associations were sourced from the Open Targets Platform (20) and the association score (y-axis) represents the aggregate evidence across all published studies for a gene-phenotype link. Only association scores >0.25 are plotted, and association scores involving the STC1 gene are highlighted with an asterisk.

To identify selective sweeps in the Turkana, we computed three statistics that rely on different assumptions and subsets of the data: 1) the integrated haplotype score (iHS) (12), computed on high coverage Turkana genomes only; 2) the population branch statistic (PBS) (3), computed on high and medium coverage Turkana genomes, and 3) the XtX statistic (13), computed on all genomes. Additionally, our PBS and XtX analyses included the Luhya (East Africa) and Yoruba (West Africa) populations from the 1000 Genomes Project as outgroups (n=207) (14). After computing all three statistics, we used a sliding window approach to intersect the results and identified 13 50kb outlier windows near 8 genes (Figure 2B, Table S6). Two of these eight genes (CCDC102B and SEMA6A) were previously found to be under selection in the Maasai, a closely related Nilotic pastoralist group (15). To our knowledge, the rest have not been previously identified as targets of selection in humans.

When we asked what phenotypes our candidate genes have been previously linked to by genome-wide association studies (GWAS), we found that they were enriched for a major predictor of cardiovascular disease—arterial stiffness. Unexpectedly, they were also enriched for neurological biomarkers of Alzheimer’s disease, namely neurofibrillary tangles, PHF-tau, and cortical surface area (Figure 2C, Table S7-8). In line with this observation, several of our candidate genes are most highly expressed in neurological cell types, such as inhibitory neurons, oligodendrocytes, and oligodendrocytes precursors (Table S9).

### Selection on STC1, a key regulator of metabolic and renal system functions

Interestingly, some of the strongest evidence from previous GWAS localized to STC1, a gene that encodes a glycoprotein with autocrine and paracrine functions. Specifically, previous GWAS have linked STC1 to 1) serum levels of urate, a waste product produced when the body breaks down purine-rich foods like red meat, and 2) serum levels of urea and creatinine, two common biomarkers of kidney function (Figure 2C, Table S7). Beyond GWAS, model organism work has implicated STC1 in glucose homeostasis (16) as well as the response to dehydration. In particular, STC1 transcription is induced up to 8-fold in rodent kidneys following water deprivation (17), a response that is coordinated by antidiuretic hormone (ADH) (18) and involves STC1 regulation of both rising hypertonicity and progressive hypovolemia (19). Given the clear involvement of STC1 in metabolic and renal system traits of ecological relevance to the Turkana, we prioritized this gene for follow-up analyses.

We identified two overlapping 50 kb candidate regions near STC1, with the collapsed 75 kb region located ∼150 kb upstream of the transcription start site in a putative enhancer. This region was ranked first out of all tested regions by the iHS statistic and third by the PBS and XtX statistics (Table S6). Previously published HiC data from five tissues that express STC1 (21) show that this regulatory region and the STC1 gene body fall within the same topological domain (22) and are in consistent contact (23) (Table S10, Figure S11). In our main tissue of interest, the kidney, STC1 is highly cell-type specific and is expressed almost exclusively in the collecting duct (24, 25). This structure is the final segment of the kidney to control fluid balance, accounting for ∼5% of water reabsorption at baseline and up to ∼25% during ADH surges induced by dehydration. When we analyzed ATAC-seq data previously generated from a mouse collecting duct cell line exposed versus unexposed to ADH (26), we found that our candidate region contained several differentially accessible, ADH-responsive regulatory elements (Figure 3A). While there is, unfortunately, no human collecting duct cell line, we performed new experiments to show that STC1 is induced by ADH in a human kidney (epithelial) cell line (linear model p-value for 10nM and 25nM, respectively = 0.033 and 0.0042; Figure 3B, Table S11).

**Figure 3.**
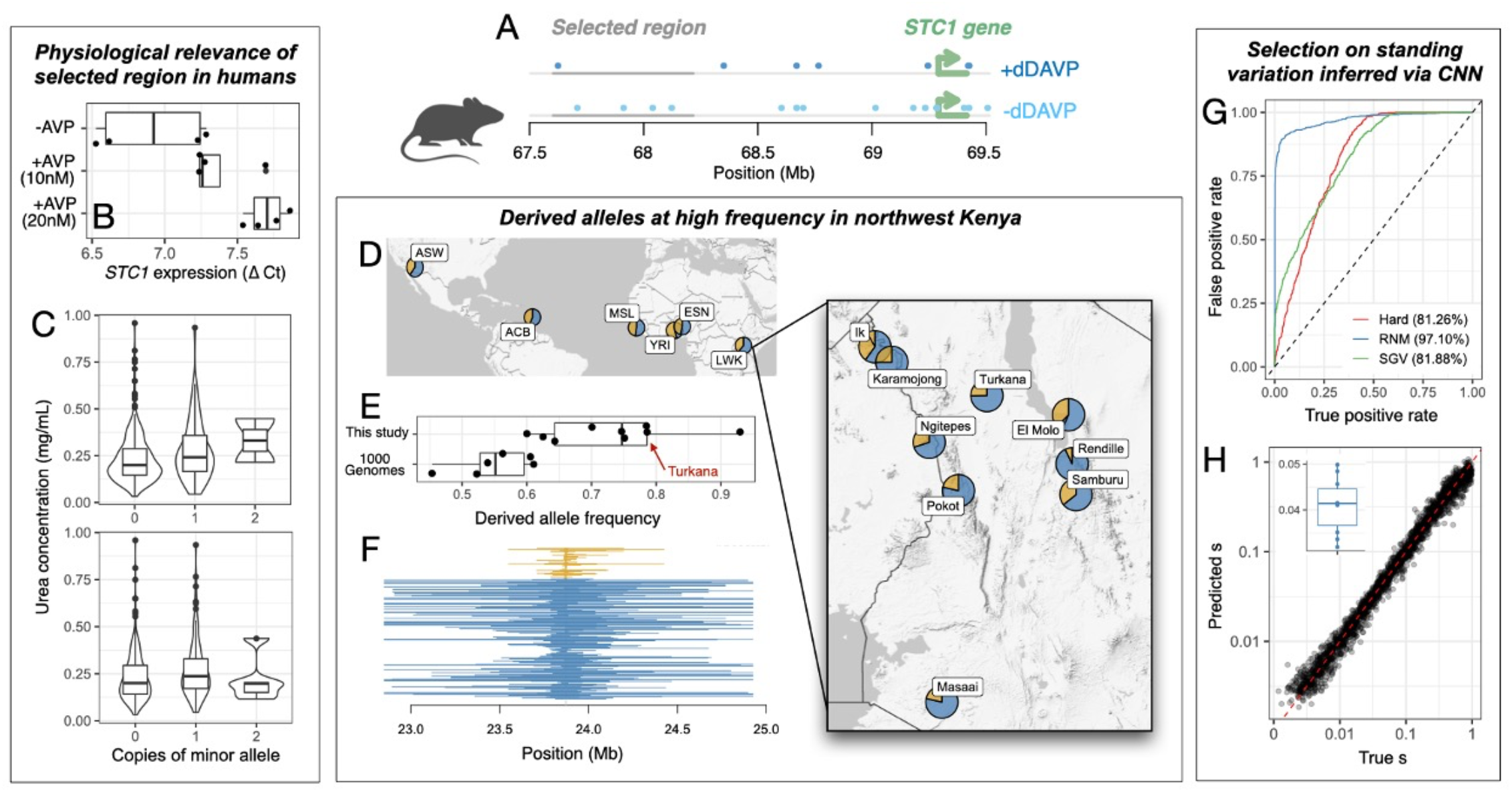
Selection near the STC1 gene. A) Open chromatin regions in mouse collecting duct cells (mpkCCD) exposed versus unexposed to ADH (26). Coordinates are in mm10; open chromatin regions are highlighted in blue and the STC1 candidate region is highlighted in grey. B) qPCR of STC1 expression levels in human kidney cells (HEK-293) exposed versus unexposed to ADH. C) Correlations between Turkana genotypes within the STC1 region and serum urea levels (FDR<5%). D) Allele frequency of the candidate region tag SNP (rs6988698) with the strongest evidence for selection. World map shows the frequency of the major (blue) versus minor (yellow) alleles in all African populations included in the 1000 Genomes Study (14); inset shows the frequency for all populations sampled as part of this study. E) Boxplot summarizing the frequency of the rs6988698 derived (major) allele. F) Haplotype length at the rs6988698 SNP for individuals carrying the derived (blue) versus ancestral (yellow) allele. Coordinates are in hg19. G) Validation of the convolutional neural network (CNN) used to infer sweep mode. ROC curves show the results of applying one CNN replicate on the simulated validation dataset. Each curve represents a one-versus-all comparison between the reference mode and all others combined; the area under each curve is given in parentheses. H) Validation and application of the CNN used to infer selection coefficients (s). Scatterplot shows the results of applying one CNN replicate on the simulated validation dataset, with the expected and predicted values of s on the x- and y-axes, respectively. Inset shows the results of applying 10 replicate CNNs to real data from the STC1 locus, under the assumption that the selective sweep is codominant (see Figure S15 for results when the sweep is assumed to be dominant).

To further link our candidate region to kidney function, we measured serum creatinine and urea levels for 447 Turkana included in our genotyping array dataset. After subsetting to SNPs within the STC1 candidate region that passed our filters, we were able to test 6 SNPs for associations with creatinine, urea, and glomerular filtration rate (estimated from the creatinine data); we note that SNP density in the array dataset is generally sparse, and these 6 SNPs are thus expected to tag general haplotype structure rather than to assist with fine mapping. Using this approach, we found 2 SNPs that were significantly correlated with urea levels (linear mixed effects model; rs10107949 beta=0.030 and p=1.96×10-2, rs75070347 beta=0.040 and p=9.23×10-3; Figure 3C, Table S12-13). While further work is needed to identify the causal variants in the STC1 region and their precise mechanism of action, these data, in aggregate, point toward selection on variants associated with STC1 regulation in the context of ADH induction, dehydration stress, and kidney function.

### Evolutionary history of STC1

We next turned our attention toward understanding the evolutionary history of the *STC1* regulatory region, including the nature, strength, and timing of selection. We started by checking the worldwide allele frequencies of the top three derived SNPs with the strongest evidence for selection in the WGS dataset (rs6988698, rs7012892, rs6994711). We found that the same SNPs that are at high frequency in the Turkana (with allele frequencies of 75%, 84%, and 84%, respectively) are near invariant outside of Africa, with average minor allele frequencies of 2.66% (range=0-8.65%; Figure S12 and Table S14). These three example variants are at intermediate frequencies in non-Nilotic East African groups, and at Turkana-like frequencies in the other East African groups we sampled (Figure 3D-F). These East African groups are almost all part of the same Nilotic lineage the Turkana belong to, which began pastoralist practices ∼5-8 thousand years ago (5, 6). Thus, we hypothesize 1) genetic variation at this region was lost during the out-of-Africa migration and 2) there was selection on standing variation within the Nilotic cluster starting sometime after pastoralist practices emerged in this group.

To test this scenario, we estimated the site frequency spectrum for the Turkana and inferred their demographic history (Figure S13-14, Table S15). We then simulated genomic datasets under different evolutionary scenarios—varying the nature, strength, and timing of selection in the *STC1* region—and trained a convolutional neural network to infer these parameters (27) (Figure S15). When we applied the trained neural network to our real data, it estimated that selection on standing variation began ∼348 generations ago (range 187-571). This timing corresponds well with the end of the African Humid Period when East Africa began to experience marked aridification (28), as well as when archaeological evidence indicates that pastoralism first emerged in the region (29, 30). Selection on the *STC1* region was inferred to be quite strong, with a selection coefficient of ∼0.041 (range 0.033-0.05; Figure 3G-H, Figure S16), in line with some of the most well-known examples in recent human history. For example, selection coefficient estimates for the malaria-protective Duffy (31) and lactase persistence (32, 33) alleles range from 0.04-0.08 and 0.04-0.06, respectively.

### Polygenic selection on metabolic and renal system biomarkers

While selective sweeps—such as the one we have detected near STC1—are undoubtedly important in human evolution (2), complex traits are more commonly shaped by weak but simultaneous selection on numerous genetic variants (34). We were therefore motivated to also explore the contribution of so-called polygenic adaptation to Turkana’s metabolic and renal system traits. To do so, we collapsed our three selection statistics into a Fisher’s combined score (FCS) and asked whether regions previously associated with 29 biomarkers (via multi-ancestry GWAS in the UK Biobank) exhibited higher mean FCS scores in the Turkana than expected by chance (35). These analyses revealed robust evidence of polygenic selection on metabolic biomarkers, such as triglycerides, cholesterol, and HbA1c, as well as renal system biomarkers, such as uric acid and cystatin C (all FDR<5%; Figure 4A, Table S15). These results suggest that there has been diffuse, genome-wide selection on metabolism and kidney function in the Turkana, in addition to the localized signal we uncovered at STC1.

**Figure 4.**
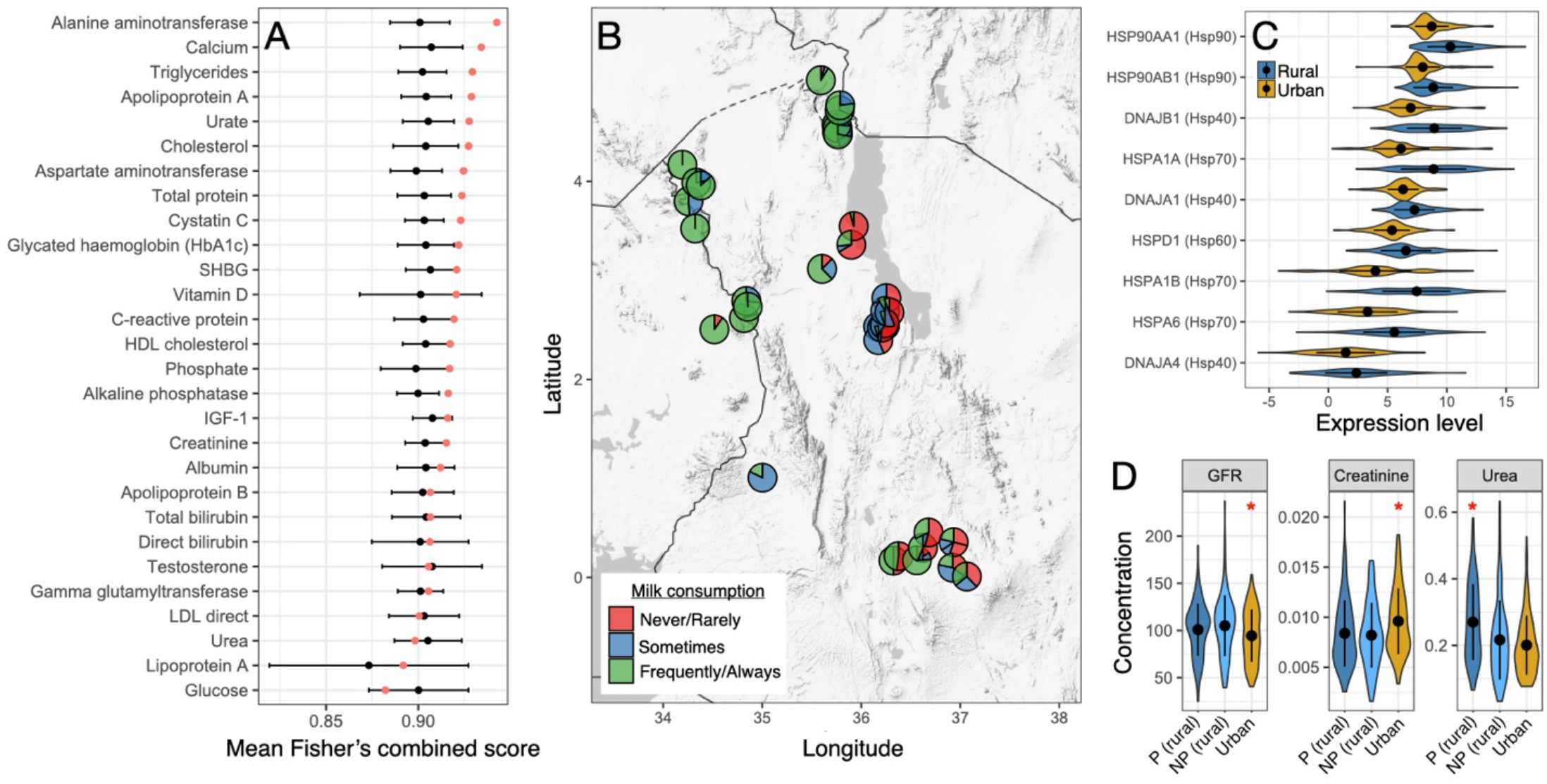
Polygenic selection and lifestyle change. A) Red dots represent the mean Fisher’s combined score (representing a composite of all three selection statistics) for each 50 kb window that contained a SNP associated with the focal trait (associations were sourced from multi-ancestry GWAS in the UK Biobank). Boxplots show the FCS distribution for 10,000 randomly sampled windows matched for SNP density, recombination rate, and background selection. B) Self-reported milk consumption summarized by location (for Turkana individuals only). C) Normalized gene expression levels for samples collected from Turkana individuals in rural versus urban locations. Because of small sample sizes, the rural category includes pastoralist and non-pastoralist individuals (see (36)); we note that results are qualitatively similar when they are split (Figure S18). Plot includes all heat shock proteins in the “response to heat” gene ontology category that were significantly differentially expressed at a 5% FDR. D) Serum concentrations of urea (mg/mL) and creatinine (mg/mL), as well as estimated GFR (mL/min), for rural pastoralists (P), rural non-pastoralists (NP), and urban individuals. Red asterisks indicate groups that were significantly different from the other two (FDR<5%).

### Lifestyle change and the fate of selected alleles

While many Turkana still practice nomadic pastoralism in northwest Kenya, and thus experience the ecological selection pressures we have highlighted here, subsets of the population are undergoing different types and degrees market integration, urbanization, and acculturation. In recent decades, some Turkanas have stopped practicing pastoralism, but continue to live relatively subsistence-level lifestyles in northwest Kenya, while others have migrated to urban areas in favor of wage labor jobs. Not surprisingly, moving to a city represents a radical environmental shift: for example, while >90% of pastoralists regularly consume blood, milk, and red meat, these numbers drop to 0%, 47%, and 31%, in urban areas where large portions of the diet are instead derived from processed foods (36, 37) (Figure 4B).

This ongoing and dramatic lifestyle shift is relevant to our evolutionary genomic analyses, because it sets the Turkana up for “evolutionary mismatch” (38). This occurs when previously advantageous variants, selected for in past ecologies, are placed in novel environments where they instead have detrimental effects. In support of this idea, we have previously shown that transitions from pastoralist to urban lifestyles are associated with increases in biomarkers of cardiovascular disease risk (36). Here, we generated new data on 1) serum urea and creatinine levels, and found that both kidney function biomarkers also differ between urban and rural/pastoralist Turkana (n=447, all FDR<5%; Figure 4D, Table S13) as well as 2) blood gene expression levels, and found that genes involved in biological processes such as “response to heat”, “protein folding”, and “inflammatory response” are differentially regulated between urban and rural/pastoralist communities (n=230, all FDR<5%; Figure 4C, Figure S17, Table S17-18). Further, we found that SNPs that fall near genes differentially expressed by lifestyle exhibit more evidence for selection than SNPs near genes unaffected by lifestyle change (linear model p-values: iHS<2×10-16, PBS=0.007, XtX=0.0211; Table S19). This result suggests that past adaptations are poised to interact with the environmental and physiological shifts the Turkana are currently experiencing.

## CONCLUSIONS

We integrated anthropological, physiological, and genomic datasets to show that an arid ecology combined with pastoralist practices has led to selection on STC1 and renal and metabolic systems more broadly in the Turkana. While we do not know the causal/selected allele at this time, we show that the STC1 gene is induced by ADH in human cells, that STC1 variants are linked to urea levels in the Turkana themselves, and that strong selection occurred on standing variation near STC1 on an ecologically plausible timescale. We also shed rare empirical light on the popular evolutionary mismatch hypothesis, which is commonly invoked to explain the high rates of non-communicable, “lifestyle” diseases observed in Western countries (39–41), but for which few appropriate study systems are available. Our work highlights that partnerships with subsistence-level groups can lead to new models for understanding how present-day environments interact with past adaptations to influence disease risk.

## ACKNOWLEDGMENTS

This work was funded by internal awards from Princeton University to JFA, The John Templeton Foundation (grant #48952 to SM), and The National Institute of General Medical Sciences of the National Institutes of Health (R35GM124827 to MAW). AMTO was supported by The Graduate College at Arizona State University, The Achievement Rewards for College Scientists Foundation Phoenix Chapter as a Pierson Scholar, and Arizona State University Sigma Xi. AJL was supported by the Helen Hay Whitney Foundation. The authors acknowledge Research Computing at Arizona State University for providing high-performance computing resources that have contributed to the research results. The National Museums of Kenya provided institutional support to conduct the research in Kenya. We thank our ASU-team research assistants for aiding with data collection: E. Carlystus, A. Lotiira, C. Muya, G. Topos, and D. Lomelu. We thank previous members of the THGP for their contributions, especially S. Lowasa, D. Mukhongo, S. Ngatia, E. Loowoth, and M. Ndegwa. We are also grateful to the staff of Mpala Research Centre for their essential support, especially F. Hassan, C. Nzomo, B. Wanjohi, G. Chege, and T. Maina. We thank Ian Wallace for thoughtful comments on a previous version. Above all else, we thank our participants and host communities for their hospitality and for their continued support in this project.

## ETHICS APPROVAL

From the inception of this project, we have engaged local area chiefs, community elders, and health officials at the local, county, and federal government levels. Our research efforts were guided by these conversations and data collection was overseen and conducted in collaboration with local area chiefs, community elders, and Kenyan scientists. Additional information about our community engagement procedures can be found in the Supplementary Methods. This study was approved by Princeton University’s Institutional Review Board for Human Subjects Research (IRB #10237), Maseno University’s Ethics Review Committee (MSU/DRPI/MUERC/00519/18), and Arizona State University’s Institutional Review Board for Human Subjects Research (IRB ID: STUDY00004874). We also received county-level approval for research activities, and research permits from Kenya’s National Commission for Science, Technology and Innovation (NACOSTI/P/18/46195/24671).

## Notes

### Competing Interest Statement

The authors have declared no competing interest.

