## Supplementary Methods and Figures for "Adaptations to water stress and pastoralism in the Turkana of northwest Kenya"

The supplement consists of:

***Supplementary Methods***

***Supplementary Figures 1-18***


- **Figure S1.** Map of current homelands/major areas for populations included in this study.
- **Figure S2**. Imputation accuracy in low/medium coverage whole genome sequencing (WGS) data is higher with a one panel approach.
- **Figure S3**. Principal components analysis of whole genome sequencing data (focusing on samples included in this study).
- **Figure S4.** Genetic variation does not systematically differ by sequencing location or coverage.
- **Figure S5.** Principal components analysis of array genotype data.
- **Figure S6.** Ancestry group and geography predict genetic variation.
- **Figure S7.** Cross-validation error (CV) obtained from the program ADMIXTURE run with different values of K. A
- **Figure S8.** Global ancestry analysis with ADMIXTURE.
- **Figure S9**. Low levels of European ancestry in the Turkana.
- **Figure S10.** Principal components analysis of whole genome sequencing data (combined with data from the 1000 Genomes Project).
- **Figure S11**. HiC data from tissues that robustly express STC1.
- **Figure S12**. Minimal genetic variation in the STC1 candidate region in Europeans.
- **Figure S13**. Overview of demographic scenarios explored for the Turkana population.
- **Figure S14**. Site frequency spectra calculated from the empirical data versus under different demographic scenarios.
- **Figure S15.** Using a convolutional neural network (CNN) to infer the mode, strength, and timing of selection.
- **Figure S16.** Results of applying CLUES to SNPs within the STC1 candidate region.
- **Figure S17**. Biological processes enriched within the set of genes differentially expressed by lifestyle.
- **Figure S18**. Biological processes enriched within the set of genes differentially expressed by lifestyle.


***Supplementary Tables 1-19***


- **Table S1**. Summaries of interviews and urine specific gravity data collected from Turkana pastoralists
- **Table S2.** Metadata for whole genome sequencing data generated by the Turkana Health and Genomics Project
- **Table S3.** Metadata for whole genome sequencing data generated by the ASU team
- **Table S4.** Metadata for the Infinium Global Screening array dataset
- **Table S5**. Results from a linear model predicting ADMIXTURE ancestry components as a function of coverage and sequencing group (for Turkana individuals only)
- **Table S6.** Candidate regions for selective sweeps (coordinates are in hg19)
- **Table S7.** Phenotypic associations from Open Targets Platform for all candidate genes

- **Table S8.** Results from Fisher's Exact Tests asking whether the candidate gene set is enriched for associations with certain traits
- **Table S9**. Cell type-specific expression information for each candidate gene (sourced from The Human Protein Atlas)
- **Table S10.** Topological domains overlapping the STC1 candidate regulatory region, for 5 tissues that robustly express STC1 (coordinates are in hg19)
- **Table S11**. STC1 gene expression levels in human kidney cells treated versus untreated with vasopressin (AVP)
- **Table S12**. Metadata for samples used to measure kidney function biomarkers
- **Table S13**. Results from linear mixed effects models of variation in kidney function biomarker levels
- **Table S14**. Allele frequency data for rs6988698 (the example tag SNP plotted in Figure 3)
- **Table S15**. Results from demographic inference performed with DaDi
- **Table S16**. Results from analyses testing for polygenic selection on metabolic and renal system traits
- **Table S17**. Metadata for samples used to measure gene expression levels
- **Table S18.** Results from gene set enrichment analyses performed on genes differentially expressed by lifestyle (rural versus urban)
- **Table S19.** Results from a linear model testing for differences in selection signal between differentially expressed (DE) and non-DE genes; all models control for gene expression level

***Supplementary References***

### SUPPLEMENTARY METHODS

***Overview of the study populations***

The Turkana are part of the Eastern Nilotic lineage; the ancestors of the present day Turkana likely entered Kenya in the early 18th century, and expanded to their current range (now known as Turkana county) by ~1900 (*1*). Today, the Turkana represent the second largest pastoralist group in Kenya after the Maasai. Turkana county is a semi-arid desert characterized by low annual rainfall, frequent droughts, and high year round temperatures (Figure 1D) (*2*).

The Turkana are traditionally nomadic pastoralists; they herd five species of livestock (dromedary camels, zebu cattle, fat tailed sheep, goats, and donkeys) and rely on these animals for subsistence (*3*). As a result of their pastoralist lifestyle, the Turkana have a remarkably protein-rich diet: 62% of calories are derived from milk or milk products, and 70-80% of calories are derived from animal products of some sort (*3*). Daily protein intake exceeds the FAO/WHO requirements by >300%, despite total caloric intake being limited (1,300–1,600 kcal/day) (*4*). Dietary items not derived from livestock are obtained through foraging or trade. For detailed descriptions of the diet, climate, and lifestyle experienced by traditional, pastoralist Turkana, see work from the South Turkana Ecosystem Project (summarized in (*5*)).

While this study focused on the evolution and present-day phenotypic variation of the Turkana people, we also worked with, and generated genomic data from, select groups that live in or near the Turkana region and/or are thought to have a shared history with the Turkana (Figure S1). This additional data generation was done to situate the Turkana within the broader context of the region and to understand population genetic parameters that could impact our selection analyses. Specifically, we generated genomic data from: El Molo, Ik, Karamojong, Masaai, Ngitepes, Pokot, Rendille, and Samburu. El Molo and Rendille are part of the Cushtic lineage and speak Afroasiatic languages, while the remaining groups are part of the Nilotic lineage and speak Nilo-Saharan languages. In recent history at least, the main subsistence strategy of the El Molo is fishing, while the Ik are agriculturalists. Like the Turkana, the Masaai, Pokot, Samburu, Rendille, Ngitepes, and Karamojong practice pastoralism. Previous work has shown some genetic substructuring by language family as well as geography for groups inhabiting northwest Kenya and northeastern Uganda (including but not limited to the groups named here) (*6*, *7*). However, none of this previous work has included whole genome sequence data and/or this entire collection of groups with geographic, cultural, or potential evolutionary relationships to the Turkana.

***Overview of community engagement and study teams***

Data and samples reported in this study were collected by two groups: 1) the Turkana Health and Genomics Project (co-directed by JFA, AJL, DM, JK, and including KMG, JE, PK, FL, AL, SL, CM, BM; turkanahgp.com) and 2) a research team based out of Arizona State University that includes AMTO, RS, CH, ACS, SM, and MAW. The Turkana Health and Genomics Project (THGP) is an ongoing, long-term study focused on the interacting roles of genotype and rapid lifestyle change in the development of non-communicable disease in the Turkana people. It is an international, multidisciplinary collaboration involving principal investigators from the US (JFA, AJL) and Kenya (DM, JK), a field team made up of Kenyan (including Turkana) scientists and technicians, and Turkana communities spanning northwest and central Kenya. Consistent with previous recommendations and ethical best practices (*8–10*), before initiating the THGP as well as this project specifically, we engaged in several years of community consultations with local stakeholders. These consultations involved working with local Turkana chiefs and elders, health workers, local scientists, and potential Turkana participants to develop our recruitment and sampling protocols.

As part of the THGP, interviews, biological samples, and phenotypic measures for this study were collected in Turkana, Laikipia, and Trans-Nzoia counties between April 2018 and February 2020.

During this time, members of the THGP visited locations where Turkana and other groups of interest were known to reside. At each sampling location, local chiefs and elders were first consulted about the project. If they believed the study to be of interest to their community, a larger meeting was held to explain the project to all interested individuals. After this period of discussion, adults (>18 years old) of self-reported Turkana identity (or other identities where appropriate) were invited to participate in the study. The study involved a structured interview, blood and urine sample collection, and anthropometric measurements as described in detail in previous work (*11*, *12*).

Saliva and cheek swab samples were collected by the ASU team from adults (>18 years old) of self-reported Turkana identity between October 2016 and 2017 in Turkana county, as described in (*13*). Both SM and CH have worked in this region for over a decade and have established and maintained a strong relationship with local communities. Community engagement by the ASU team proceeded in a very similar way as for the THGP, with conversations starting at the community level and funneling down to the individual level (*13*).

More specifically, for ASU-led as well as THGP-led data and sample collection, we worked with team members that were fluent in local languages (e.g., Turkana, Kiswahili) to assist with initial community-level explanations of the project as well as informed consent. In all cases, team members discussed the study with each potential participant and provided ample time for questions and concerns to be voiced. Formal consent was then obtained, and all individuals who agreed to take part in the study were provided with the contact information for both local and foreign research team members in the event that s/he should choose to be removed from the study. Throughout data and sample collection, we worked to ensure ethical informed consent and multiple levels of communication about the study. All data and sample collection was conducted in collaboration with individuals from the participant communities, many of whom have worked with our various scientific teams for many years. We have also made an effort to share our results with study communities, including the results of this study (see turkanahgp.com for information on results sharing by the THGP). The ASU team designed culturally appropriate images in consultation with local field assistants and returned results back to communities in 2018 (AMTO) and 2022 (RS, SM), with further dissemination efforts planned for 2023.

Relevant scripts:

- F_map_samples.R

***Interview data collection***

Structured interviews were conducted with all THGP participants as described previously (*11*, *12*). All interviews were conducted in a language familiar to the participant (English, Turkana, or Kiswahili). The following self-reported variables from the interviews were used in this study: 1) age and gender; 2) main subsistence activity, chosen from the following categories: self-employment, formal employment, petty trade, farming, pastoralism, hunting and gathering, other; 3) how much of the day was spent finding and retrieving water, chosen from the following categories: less than 1 hour, 1 hour, a few hours, half the day, most of the day; and 4) whether s/he would drink more water if they had it.

We also used a food frequency questionnaire to collect information about the consumption of blood, meat, milk, bread, sugar, salt, and cooking oil. We focused on these items because they reflect foods that are essential (blood, meat, milk) or uncommon (bread, sugar, salt, cooking oil) in the diet of traditional pastoralists. Participants were asked how often a specific item was used or consumed and were given the following answer choices: never, rarely, 1–2 times per week, >2 times per week or every day. These interviews were conducted year round and results therefore summarize over known seasonal variation in dietary patterns among Turkana pastoralists.

***Genomic data generation***

From THGP-collected biospecimens, we generated 1) high coverage (>200 million paired end reads) whole genome sequencing (WGS) data for Turkana individuals only; 2) medium coverage WGS data for all 9 groups (Turkana, El Molo, Ik, Karamojong, Masaai, Tepes, Pokot, Rendille, and Samburu); 3) array genotype data for all 9 groups; and 4) transcriptomic data for Turkana individuals only (described in the next section) (Table S2, Table S4, and Table S16). From ASU-collected biospecimens, we generated high coverage whole genome sequencing (WGS) data for Turkana individuals only (Table S3). The specific biospecimens and laboratory procedures used to generate these data are described below.

Genomic data for THGP samples was generated from venous blood, while genomic data for ASU samples were generated from saliva . Participants were asked to rinse their mouth with water if they recently had been chewing tobacco or other organic products; study team members then watched and guided participants to collect saliva using appropriate technique. Saliva samples were collected using the Oragene OG-500 DNA collection kit. All samples were stored short term in a cooler bag with frozen ice packs before they could be stored long-term at -4C. DNA was extracted and prepared for WGS using the Promega ReliaPrep kit and sequencing was performed on the Illumina NovaSeq S4 platform using paired-end, 2x150 reads; DNA extraction and sequencing were performed at the Yale Center for Genomic Analysis. Information on the tissue source, sequencing batch, read depth, and other metadata for each ASU WGS sample can be found in Table S3.

For THGP samples, approximately 6mL of intravenous blood was collected into an EDTA vacutainer tube. Whole blood was then mixed with DNAGard (Sigma) following the manufacturers' instructions and stored in a cooler box with frozen ice packs until the sample could be stored long-term at -20C. Samples were later thawed and 200ul of each sample was used to extract genomic DNA using the Zymo Quick-DNA Minprep kit following the manufacturers' instructions. All DNA samples for high coverage WGS were prepared using either the NEBNext Ultra DNA Library Prep Kit for Illumina (New England BioLabs) or the Nextera DNA Flex Library Prep Kit (Illumina). In both cases, we followed the manufacturers' instructions, but when using the Nextera kit, we scaled reactions to ¼ volumes.

A subset of libraries for low coverage WGS were prepared as outlined in (*14*), using the oligo sequences and homemade Tn5 recipe therein. Specifically, 10µl (100µM) forward oligo adapter A and 10µl (100µM) reverse oligo (Tn5MErev) were mixed with 80µl reassociation buffer (10mM Tris pH 8.0, 50mM NaCl, 1mM EDTA), and annealed in a thermocycler following the program: 95°C 10min, 90°C 1min, reduce temperature by 1°C/cycle for 60 cycles, hold at 4°C. The same procedure was also performed using forward oligo adapter B. To load the pre-annealed adapters onto the Tn5, 45µl of Tn5 was combined with 9µl of pre-annealed adapter A and 9µl of pre-annealed adapter B (both at 10µM) and incubated in a thermal cycler for 30min at 37°C. The pre-charged Tn5 was then diluted with reassociation buffer:glycerol (1:1), in a ratio of 1 part reassociation buffer:glycerol to 1 part pre-charged Tn5. 2µl of each DNA sample was then mixed with 1µl of precharged Tn5, 2µl of 5X TAPS buffer pH 8.5 (50mM TAPS, 25mM MgCl2, 50% v/v DMF), and 5µl of water. The solution was incubated for 7min at 55°C, after which 2.5µl of 0.2% SDS (Promega, #V6551) was added and incubated in a thermal cycler for 7min at 55°C to dissociate the Tn5 from the DNA. Final library amplification was then performed by combining 2µl of the tagmentation reaction with 7µl of OneTaq HS Quick-Load 2x (NEB, #M0486L), 1µl of the i5 primer (5uM), 1µl of the i7 primer (5µM), and 4µl of water. This mixture was amplified on a thermal cycler with the following program: 68°C 3min, 95°C 30sec, [95°C 10sec, 55°C 30sec, 68°C 30sec] for 12 cycles, 68°C 5min. All THGP WGS libraries were sequenced on the Illumina NovaSeq S4 platform using paired-end, 2x150 reads at the Genomic Core Facility at Princeton's Lewis Sigler Institute for Integrative Genomics. Information on the sequencing batch, read depth, and other metadata for each THGP WGS library can be found in Table S2.

In addition to WGS data generation, a set of 994 THGP DNA samples were also sent for SNP genotyping on the Infinum Global Screening Array at the Children's Hospital of Philadelphia (CHOP).

***Low level processing of array genotype data***

We genotyped a total of 994 samples across 9 batches. Genotype clustering was performed by CHOP using Illumina's GenomeStudio. Genotypes were obtained from CHOP and converted to Plink map format (forward strand) using Illumina's GenomeStudio and the Plink plugin. We merged the genotypes from each batch together using Plink and then filtered for autosomal SNPs with MAF>1%, SNPs in Hardy Weinberg equilibrium ($p>10^{-6}$), SNPs called in >95% of samples, and SNPs not in LD. Specifically, we used the indep-pairwise function in Plink to scan windows of 50kb with a 20kb offset, and to randomly prune variants within each window so that no pair exceeded an $R^2$ threshold of 0.8 (as in (*15*)). This filtering resulted in a dataset of 329,197 SNPs, with 400 SNPs failing our Hardy Weinberg filter, no SNPs failing our missingness filter, 219,587 SNPs failing our MAF filter, and 23,419 SNPs in LD removed. We used these 329,197 remaining SNPs as the input for KING (*16*) to estimate IBD and to prune pairs of second-degree relatives (IBD>0.125) from the dataset. We also used KING to estimate the concordance of heterozygous genotypes for 21 samples that were genotyped twice across batches to assess reproducibility, which we found to be very high (mean heterozygous genotype concordance = 1.000). After removing close relatives, duplicate samples, and 6 samples for which the Plink-infered sex (using the sex-check function) did not match the expected sex from our interview data, we were left with 782 samples (Table S4).

We used functions within Plink to perform PCA on this filtered sample set (Figure S5). We then used linear models to estimate the contribution of self-identified ancestry group to overall genetic variation, as well as the contribution of geography (longitude and latitude) to genetic variation within the Turkana. In particular, we used linear models to estimate the percent variance explained ($R^2$) by group or longitude and latitude for each of the top 10 PCs (Figure S6).

Relevant scripts

- C1_process_QC_PCA.sh

***Low level processing of WGS data***

We sequenced a total of 110 high coverage samples and 350 low and medium coverage samples. Following sequencing, we removed adapter contamination and low-quality bases from each library using cutadapt (*17*). We mapped the trimmed reads to the GRCh38 version of the human reference genome using BWA (*18*), and used Samtools (*19*) to retain only uniquely mapped reads. On average, $94.33 \pm 0.63\%$ and $88.12 \pm 4.36\%$ of our reads mapped uniquely for blood-derived and saliva-derived samples, respectively. We then marked duplicates and performed base quality score recalibration, application, and individual variant calling following the Genome Analysis Toolkit best practices pipeline (*20*). More specifically, we 1) sorted the trimmed, mapped reads using Samtools (*19*); 2) we used Picard's MarkDuplicates function (*21*) to identify duplicate reads and add read groups; 3) we performed base quality score recalibration and application using the BaseRecalibrator and ApplyBQSR functions within GATK (using the 1000 Genomes phase 3 calls as the known variants file); and 4) we created individual GVCF files using HaplotypeCaller function within GATK (v4.1.4). In cases where a given sample was sequenced across multiple lanes and therefore had multiple FASTQ files, we processed these files independently for trimming, mapping, and filtering, and then combined all of the bam files for a given individual at the MarkDuplicates step.

Relevant scripts:

- G1_trim_map.sh
- G2_bam_processing.sh

***Joint genotyping and phasing of WGS data***

We performed two rounds of joint genotyping using 1) only the 110 high coverage samples and 2) only the 350 low and medium coverage samples. In both cases, we followed the GATK best practices guidelines, such that samples were first combined using the CombineGVCFs function and then jointly called using the GenotypeGVCFs function. Using the jointly called high coverage genotypes only, we followed GATK's guidelines for filtering and final variant discovery. We later selected these sites from the jointly called low coverage genotype call set and then imputed any missing data using the high coverage call set as a reference. Our specific procedures and pipelines are detailed below. In this section, we focus on the filtering, variant discovery, and phasing of the high coverage call set; in the next section, we discuss how these filtered variants were identified and imputed in the low and medium coverage samples.

Following joint genotyping of the high coverage samples, we filtered to remove the following loci that were not appropriate for our analyses:

- Indels and deletions
- Non autosomal SNPs
- SNPs not in Hardy Weinberg equilibrium ($p<10^{-6}$; threshold as in (*22*))
- SNPs with a MAF<0.01
- Sites within segmental duplications as defined by: http://hgdownload.cse.ucsc.edu/goldenPath/hg38/database/genomicSuperDups.txt.gz
- Sites within a CpG site
- Sites within the 1000 Genomes accessibility mask as defined by: http://ftp.1000genomes.ebi.ac.uk/vol1/ftp/data_collections/1000_genomes_project/working/20160622_genome_mask_GRCh38/PilotMask/

This filtering resulted in us retaining the following numbers of loci:

- 28,143,508 out of 32,818,930 total called variants were SNPs (and not indels/deletions)
- 19,801,886 SNPs passed our region filters (i.e., were not within segmental duplications, the 1000 Genomes accessibility mask, or CpG sites)
- 19,724,008 variants passed our Hardy-Weinberg filters (77,878 were removed)
- 12,243,197 SNPs pass our MAF filters (7,480,811 were removed)

Using this set of 12,243,197 SNPs, we recalibrated variant quality scores using GATK's VariantRecalibrator function. As recommended by the GATK best practices pipeline, we used the 1000 Genomes phase 1, HapMap, and 1000 Genomes Omni array datasets as reference databases. Following variant recalibration, we filtered based on the algorithm estimated sensitivity-specificity thresholds using a tranche of 90.0 (relatively conservative). Filtered variants were then lifted over to hg19 for phasing using Picard's LiftoverVcf function (note that this liftover resulted in a loss of ~0.06% of variants) (*21*). We performed phasing on all biallelic sites with <5% missing data across all samples using SHAPEIT2 (*23*) and the 1000 Genomes phase 3 genetic map. These procedures resulted in a total of 7,767,165 phased genotypes.

Relevant scripts:

- G3_joint_call_high_cov.sh
- G3_joint_call_low_cov.sh
- G4_filter_high_cov.sh
- G5_phase_high_cov.sh

***Imputation of low and medium coverage WGS data***

Following joint genotyping of the low and medium coverage samples, we filtered for the 7,767,165 sites passing filters in our phased, high coverage call set. We also removed samples that did not have genotype quality scores >10 at >5% of all loci we were interested in. This resulted in removal of 20 samples. For the remaining 330 samples, we masked all genotype calls with genotype quality scores <10 and used this call set as the input for imputation. Prior to imputation, we used Picard's LiftoverVcf function to transfer our site coordinates to hg19.

Following previous examples in the literature (*22*, *24*, *25*), we tested two imputation strategies using IMPUTE2 (*26*). Specifically, we assessed the accuracy of imputation using 1) our phased reference panel of 110 high coverage Turkana genomes or 2) a phased reference panel of the high coverage Turkana genomes as well as a second phased reference panel of 1000G phase 3 variants. In case of such panel combining, IMPUTE2 imputes only genotypes for variants that are present in the first (main) panel, but in the process also draws on haplotype information from the second panel to potentially improve imputation accuracy. In both pipelines, low and medium coverage genomes were not "pre phased", because this increases imputation speed but potentially reduces accuracy. As recommended by the program's authors, imputation was performed on 5 megabase intervals.

The accuracy of each approach was assessed using the concordance and $R^2$ values produced through IMPUTE2's internal cross-validation procedure. More specifically, IMPUTE2 masks the genotypes from one variant at a time in the reference dataset and imputes the masked genotypes; the imputed genotypes are then compared with the original genotypes to evaluate the quality of the imputation procedure (*26*). These analyses revealed that the one panel approach produced slightly more accurate genotypes: we observed a median $R^2$ of 95.7% vs 95.1% and a median concordance of 98.1% vs 97.8% for the one panel and two panel approaches, respectively. These distributions were significantly different (Wilcoxon signed rank test; p-value for comparison of $R^2$ and concordance values=$4.88 \times 10^{-4}$ and $4.72 \times 10^{-5}$). Importantly, controlling for the proportion of genotypes that were imputed, we didn't find any significant differences in imputation performance with the one panel approach for Turkana versus non Turkana samples (linear model; beta for group comparison of $R^2$ and concordance values=$1.97 \times 10^{-3}$ and $7.51 \times 10^{-4}$, p-value=0.279 and 0.37; Figure S2). We therefore used the variant calls from this approach for all downstream analyses. We also confirmed that these imputed calls were well-correlated with genotypes from the Global Screening array, though we note that this comparison focuses on a biased set of loci that overlap between the two platforms ($R^2$ between imputed WGS and array genotypes=0.958 ± 0.025, n=108).

Relevant scripts:

- G6_filter_low_cov.sh
- G7_impute_low_cov.sh
- G8_compare_impute_array.sh
- F_assess_imputation.R

***Merging, filtering, and QC of all WGS genotypes***

Next, we proceeded to merge our high and low/medium coverage genotype call sets. Before doing so, we first filtered our imputed, low/medium coverage genotypes for well-imputed SNPs using the metrics provided by IMPUTE2. We defined well-imputed SNPs as those with $R^2$>0.8 or INFO>0.4 following (*22*), which left us with 6,476,837 SNPs. Before merging, we also removed 1) individuals that had excess levels of heterozygosity as estimated by Plink (F>0.2 or F<-0.2) (*27*); 2) duplicate samples from individuals that were genotyped via two independent libraries (retaining the library with the highest coverage; 3) 2 samples whose genotypes had been truncated due to a computational error; and 4) related individuals. For relatedness filtering, we used built-in functions in KING (*16*) to estimate IBD from the set of filtered, well-imputed SNPs and to prune pairs of second-degree relatives (IBD>0.125) from the dataset. This left us with 261 low and medium coverage samples (202 of which were from Turkana individuals) and 106 high coverage samples, which were merged using Plink (*27*).

After merging, we performed a final round of minor allele frequency filtering on the combined call set (retaining variants with MAF>0.01). We also used Plink to create an LD filtered version of the merged call set using the parameters specified in (*15*). Specifically, we used the indep-pairwise function to scan windows of 50kb with a 20kb offset, and to randomly prune variants within each window so that no pair exceeded an $R^2$ threshold of 0.8. In total, this resulted in two versions of the merged call set, one without LD filtering containing 6,355,282 SNPs and one with LD filtering containing 2,965,175 SNPs.

We used Plink to perform PCA on the LD filtered call set, and we used these principal component (PC) loadings to confirm that there were no effects of read depth or collection team/sequencing center on genetic variation captured by the first PCs. More specifically, we confirmed there were no major differences between sequencing location (Princeton vs ASU for Turkana high coverage samples) or between coverage groups (high vs low/medium for Turkana samples; Figure S3). To do so, we used Wilcoxon signed rank tests followed by multiple hypothesis testing correction and confirmed that the FDR corrected p-value for almost all tests was>0.1 (note, we observed one correlation between coverage group and PC10 at an FDR-corrected p-value of 0.068; Figure S4).

Relevant scripts:
- F_PCA_low_high_cov_data.R

***Population genomic analyses***

We merged our LD filtered dataset of 2,965,175 SNPs with the 1000 Genomes Phase 3 call set for African populations (*28*). We used Plink to filter for variants in LD (scanning windows of 50kb with a 20kb offset, and pruning so that no pairs in each window had an $R^2$>0.8)), variants with a MAF<1%, and variants genotyped in <25% of samples in the merged dataset. We then used Plink to perform PCA (Figure 2A).

We also used two approaches to understand fine scale population structure and admixture in our sample set. First, we used the program ADMIXTURE (*29*) to estimate the proportion of the genome originating from K ancestral populations for each individual, with K being specified a priori. We performed these analyses after merging our LD filtered dataset of 2,965,175 SNPs with the 1000 Genomes Phase 3 call set for the Yoruba, Luhya, and CEPH populations (*28*). We again used Plink to filter for variants in LD (scanning windows of 50kb with a 20kb offset, and pruning so that no pairs in each window had an $R^2$>0.8)), variants with a MAF<1%, and variants genotyped in <25% of samples in the merged dataset. We ran ADMIXTURE with default parameters exploring K=2-7, and each value of K was run five times with a different random seed. We chose the value of K that minimized the cross-validation error (Figure S7), which was K=3, but we also plot results when K=4 for visualization of further substructure (Figure S8). These analyses revealed small amounts of putative European ancestry within the Luhya, as was shown previously by the 1000 Genomes Project (*30*)). They also revealed small amounts of putative European ancestry within the East African groups we sampled. While the specific East African groups we worked with are mostly uncharacterized in the literature, the Simons Genome Diversity Project has reported similar ancestry breakdowns for the Masaai (*28*) and other work has confirmed admixture pulses from both Eurasia and West African into East African Afroasiatic groups (*31*). Nevertheless, to confirm that the patterns we observed were not a result of technical biases in our WGS data, we asked whether coverage or sequencing location predicted ancestry proportions within our Turkana samples, and found that this was not the case (linear model, all FDR-corrected p-values>0.05; Table S5).

We also reran the ADMIXTURE analysis with K=3 using our array data combined with array data from the 1000 Genomes Project for Yoruba, Luhya, Maasai, and CEPH (*30*). This 1000 Genomes data was generated on the Illumina Omni 2.5 array, and we therefore used Genotype Harmonizer to recover sites profiled in both datasets and mapped to the same genomic strand (*32*). We also used Plink to filter for variants in LD (scanning windows of 50kb with a 20kb offset, and pruning so that no pairs in each window had an $R^2>0.8$)), variants with a MAF<5%, and variants genotyped in <5% of samples in the merged dataset. This resulted in a dataset of 198,009 variants. When we applied ADMIXTURE to this database, we observed very similar ancestry breakdowns for all East African groups as we had seen using the WGS data (Figure S8).

Our second approach relied on local ancestry assignments from RFMix (*33*) for the high coverage samples only (because the program requires information from phased haplotypes). More specifically, RFMix attempts to assign each segment of a chromosome to its most likely ancestral source population. As potential source populations, we used CEPH individuals to represent European ancestry, Luhya individuals to represent East African (Bantu) ancestry, and Yoruba individuals to represent West African ancestry (all sourced from the 1000 Genomes phase 3 call set (*30*)). We note that this analysis does not include an East African (Afroasiatic) population as a potential source, because the sizable and publicly available datasets that include this ancestry are array rather than WGS-based (e.g., (*31*)). We also note that our main goal here was to confirm the results of the ADMIXTURE analysis (especially the European ancestry), and we therefore reasoned that it made sense to use the same populations and data. Using the RFMix method, we observe similar evidence for European ancestry as was estimated by ADMIXTURE (Figure S9). We also note that our estimates of European admixture from RFMix are highly correlated with our estimates from ADMIXTURE (Pearson correlation R=0.84, $p<10^{-16}$; Figure S9).

Relevant scripts:
- F_multi_pop_PCA.R
- A1_run_admixture.sh
- A2_run_admixture_array.sh
- A3_run_RFmix.sh

***Genome-wide scans for positive selection***

We identified genomic regions under recent positive selection in the Turkana using an outlier approach and three statistics: the integrated haplotype score (iHS) (*34*), the population branch statistic (PBS) (*35*), and the XtX statistic as computed by BayEnv2 (*36*). These approaches can detect selection on the order of thousands to tens of thousands of years ago (*37*). Specifically, the iHS compares patterns of linkage disequilibrium within a population for haplotypes carrying the derived versus ancestral SNP at a given locus, while the PBS and XtX approaches compare allele frequencies between the Turkana and two or more reference populations and look for SNPs where the genetic distance is substantially different from background. By using three statistics that each look for distinct signals in the data and rely on different aspects of the data (e.g., iHS analyzes phased haplotypes, while PBS and XtX analyze allele frequencies), we reasoned that we could recover robust signals of selection and avoid false positives induced by the processing steps or assumptions required for any individual test.

iHS relies on linkage disequilibrium information within phased haplotypes, and we therefore focused only on the high coverage genotype call set when calculating this statistic. Specifically, iHS values were calculated for the phased, high coverage samples using the R package REHH (*38*). Before running REHH, we converted the phased haplotype output from SHAPEIT2 to VCF format using the convert function in SHAPEIT2, and we added information about derived/ancestral state at each locus from 6-way EPO alignments (*39*) using the fill-aa function in VCFtools (*40*). Using the default parameters in REHH, 4,648,783 SNPs were analyzable with known ancestral state calls, data for >90%

of samples, and MAF>5%. iHS estimates were normalized internally by the program as a function of their derived allele frequency.

To calculate PBS, we started with the set of 2,965,175 imputed and LD filtered SNPs passing filters in our merged dataset of low, medium, and high coverage samples. We merged this dataset further to the Yoruba and Luhya samples from the 1000 Genomes phase 3 call set, and refiltered the merged dataset to SNPs with MAF>1%, data for >75% of samples, and LD structure that passed the same filters we used previously (as specified in (*15*)). Using this set of 2,144,319 SNPs, we calculated $F_{ST}$ in Plink for each pairwise population comparison and we then used R to calculate PBS from the $F_{ST}$ values using the equations provided in (*35*).

While BayEnv2 is most commonly used to test for associations between ecological or climatic variables and genetic differentiation, the program can also calculate an environment-agnostic population differentiation statistic called XtX. This statistic is similar to the classic $F_{ST}$ statistic and is meant to identify highly differentiated SNPs that might be driven by local adaptation. An attractive feature of XtX is that it can identify highly differentiated SNPs across an infinite number of populations (rather than just 2, in the case of $F_{ST}$, or 3, in the case of PBS), and provide information about how differentiated the focal SNP is relative to the expected, genome-wide genetic covariance between populations. To calculate XtX, we followed the workflow outlined in (*41*) and here: https://eead-csic-compbio.github.io/barley-agroclimatic-association/HOWTOXtX.html. First, we used our array data to compute a population-level genetic covariance matrix for the following groups: El Molo, Ik, Luhya (from 1000 Genomes), Pokot, Ngitepes, Karamojong, Rendille, Samburu, Turkana, and Yoruba (from 1000 Genomes). This procedure used the SNPs obtained after merging our filtered Global Screening array dataset to the 1000 Genomes Omni array call sets and refiltering for SNPs with MAF>5%, data for >95% of samples, and LD structure that passed the same filters we used previously (as specified in (*15*)). We then ran BayEnv2 for all 2,965,175 imputed and LD filtered SNPs passing filters in our merged dataset of low, medium, and high coverage WGS samples. We ran the program three times with three different random seeds and 100,000 iterations per run; we then averaged the per-site output to obtain the XtX statistics used in downstream analyses.

Relevant scripts:

- S1_run_BayEnv_part1.sh
- S1_run_BayEnv_part2.R
- S1_run_BayEnv_part3.sh
- S2_run_iHS_part1.sh
- S2_run_iHS_part2.R
- S3_run_PBS_part1.sh
- S3_run_PBS_part2.R

***Identifying candidate regions and candidate genes for positive selection***

To identify candidate regions for selective sweeps, we rank ordered the |iHS|, XtX, and PBS distributions and defined outliers as those in the top 1% of each distribution. We then binned the genome into 50 kb windows with 25kb offsets, and counted the number of outlier loci for each statistic that fell in a given window (but excluding windows with <20 SNPs tested). Windows with outlier numbers in the top 1% of the genome-wide distribution for all three selection statistics were considered as candidates. Because each statistic has its own underlying assumptions and sensitivities, windows that are outliers for many different tests are expected to be enriched for true positives (*42–44*). We required all three statistics to exhibit outliers in a given region in order to identify the most robust signals; in other words, we aimed to minimize the false positive rate even if it came at the cost of a higher false negative rate (as is common in the literature (*42–44*)). We chose to use 50 kb windows because simulation studies have shown this window size provides good power to detect sweeps with selection intensities between 2Ns=100 and 1000 (*45*). This procedure resulted in 13 50 kb windows, representing 8 non-overlapping windows (Table S6).

We considered candidate genes for positive selection to be protein-coding genes within each non-overlapping candidate window, or within 500kb of the start or end of a given candidate window. To understand the biological function of these genes, we downloaded information about transcriptional cell type specificity from the Human Protein Atlas (https://www.proteinatlas.org/). Here, we used the RNA specificity category, which is based on mRNA expression levels in scRNA-seq data from normal tissues (Table S9). To understand which human diseases and traits this set of candidate genes has been linked to, we used the Open Targets database (https://www.targetvalidation.org/). First, we looked up each candidate gene in the database and downloaded information on the traits that gene had been linked to via previous GWAS: the strength of this link is summarized via a genetic association score for each gene-trait combination (Table S7). Next, for traits associated with 2 or more genes in our dataset, we downloaded the total number of genes in the human genome associated with the trait from Open Targets. We then used a Fisher's exact test to ask whether the number of candidate genes associated with the focal trait was more than expected by chance, and we used a 10% FDR cutoff for significance (Table S8). We note that we excluded behavioral traits (e.g., educational attainment) from this analysis because of known problems and sensitivities with these phenotypes.

Relevant scripts:
- S4_summarize_outliers.R
- S5_allele_freq.sh
- F_candidate_genes.R
- F_STC1.R

***Measuring STC1's response to ADH***

Immortalized renal proximal tubule epithelial cells (RPTEC/TERT1) were purchased from ATCC (CRL-4031) and cultured according to ATCC's recommended culturing protocols. Cells were propagated and passaged, and independent experiments were performed on passages 9 and 13 (we refer to these independent experiments below as "batches"). On the day before each of the experiments, cells were trypsinized following ATCC protocols for this cell type, and plated into 6- or 12-well plates at ~70-80% confluency.

On the day of each experiment, cells were pre-treated for 20 minutes with IBMX (Cayman Chemicals 10008978) before being treating with 10 or 25 nM vasopressin (Sigma Aldrich V1005) with IBMX. For the first experiment/batch, we used PBS + IBMX as well the control and during the second experiment/batch we used PBS + DMSO as the control condition. After 24 hours of vasopressin treatment, cells were rinsed once with PBS and then lysed in RNA lysis buffer (Zymo Research R1050). RNA was purified using the Zymo Quick-RNATM Microprep Kit (R1050). We synthesized cDNA using the Applied Biosystems High-Capacity RNA-to-cDNATM Kit (ABI 4387406) and ran the qPCR using the PowerUpTM SYBR Green Master Mix (ABI A25741). We then used linear models to test for treatment effects on delta Ct values, controlling for experimental batch (treatment effect p-value for 10nM and 25nM: 0.033 and 0.0042). We did not find any evidence that batch influenced our results (batch effect p-value for 10nM and 25nM: 0.9540 and 0.12142).

***Measuring biomarkers of kidney function and analyzing sources of variance***

We used LC-MS to measure creatinine and urea from 446 Turkana serum samples that were also genotyped on the Infinium Global Screening Array and passed our quality control filters (Table S12). In addition to our Turkana samples, each run included spike-ins of both creatinine and urea to calculate standardized, absolute concentrations (CLM-311-1 UREA (13C, 99%) and DLM-1302-0.25 CREATINE (METHYL-D3, 98%)). Water-soluble metabolite measurements were obtained by running samples on the Orbitrap Exploris 480 mass spectrometer (Thermo Scientific) coupled with hydrophilic interaction chromatography (HILIC). We used an XBridge BEH Amide column (150mm X 2.1 mm, 2.5 uM particle size, Waters, Milford, MA) to prepare samples. The gradient was as follows: solvent A (95%:5% $H_2O$:acetonitrile with 20 mM ammonium acetate, 20 mM ammonium hydroxide, pH 9.4) and solvent B (100% acetonitrile) 0min,90% B; 2min,90% B; 3min,75% B;7min, 75% B; 8min,70%,9min, 70%B; 10 min, 50% B; 12 min, 50% B; 13 min, 25% B; 14 min, 25% B; 16 min, 0.5% B, 20.5 min, 0.5% B; 21 min, 90% B; 25 min, 90% B. The flow rate was set to 150 mL/min with an injection volume of 5 μL and a column temperature of 25°C. The MS scans were completed in polarity switching mode to enable both positive and negative ions across a mass range of 58–1000 m/z, with a resolution of 120,000. Data were analyzed using the EI-MAVEN software (v 0.12.0, Elucidata). Glomerular filtration rates were estimated for each individual from absolute creatinine levels using the CKD-EPI Creatinine Equation recommended by the National Kidney Foundation (https://www.kidney.org/content/ckd-epi-creatinine-equation-2021).

After performing the genotype filtering described in *Low level processing of array genotype data,* we retained 6 SNPs in the candidate region near *STC1*. For each of these SNPs, we used linear mixed effects models to test for an association between genotype (coded as 0, 1, or 2 copies of the minor allele) and each kidney function biomarker (creatinine, urea, or GFR). All models controlled for age, lifestyle, and a genetic relatedness matrix derived from the genome-wide, filtered array genotype data. Lifestyle was coded as urban, non-pastoralist (rural), or pastoralist (rural). To do so, we used the definitions in (*12*), which are briefly as follows: 1) pastoralist (rural) was defined as individuals that reported their main subsistence activity as pastoralism, that drink milk every day (i.e., that rely on their livestock for subsistence), and that live in Turkana county; 2) non-pastoralist (rural) individuals were defined as those that live in Turkana county but did not meet the criteria for category #1; and 3) non-pastoralist (urban) individuals were defined as those who no longer practice pastoralism and reside in one of three cities included in our study—Nanyuki, Lodwar and Kitale. Linear mixed effects models were implemented in the R package EMMREML, followed by multiple hypothesis testing correction using a false discovery rate approach and a 10% FDR cutoff. To understand how lifestyle influences each kidney biomarker, we also ran a set of linear mixed effects models identical to those described above but that excluded any SNP information (see Table S13).

***Inferring the demographic history of the Turkana***

We used the "include-non-variant-sites" flag in GATK's GenotypeGVCFs function to emit all callable loci in the Turkana genome (focusing on the high coverage samples only). This resulted in 2,863,768,058 autosomal loci (26,009,619 biallelic SNPs), which were further pruned to obtain high quality sites and neutral regions. To obtain high quality sites, we employed filtering based on depth (DP) and the number of alleles that were genotyped. Specifically, we first calculated the mean depth across all sites (the field DP from the VCF file). The mean DP across all sites was 259.5, and a site was considered high quality if its DP was greater than 130. In addition, a site was considered high quality if it was genotyped (the field AN from the VCF file) in at least 50% of the dataset. The filtering resulted in 2,664,308,333 autosomal loci.

To obtain neutral regions, we used the neutral explorer software (*46*). Specifically, we ran the software with the following parameters: 1) regions to exclude: known genes, segmental duplications, gene bounds, CNVs, spliced ESTs, self chain; 2) minimum region size = 200bp; 3) distance to nearest gene = 0.4cM; 4) recombination rate = 0.9cM/Mb; 5) genetic map = HapMap; 6) human diversity = Yoruba; 7) mask = strict; 8) minimum BG selection coefficient = 0.95. After filtering for loci that passed our quality filters and were in neutral regions, we were left with 44,921,784 sites (356,517 biallelic SNPs).

To infer demographic history, we ran the program DaDi (*47*) with three models (Figure S13): 1) the 1-Epoch model, which is a constant growth model; 2) the 2-Epoch model, which includes a population contraction or a population growth event; and 3) the 3-Epoch model, which includes two population contraction and/or growth events. For each model, we implemented 100 replicates. To visualize the fit of the different models to the empirical data, we used the inferred parameters to generate the site-frequency-spectrum.

We found that the constant size model (1-Epoch) resulted in the worst likelihood (Table S15) and the site-frequency-spectrum generated using the parameters inferred from this model did not fit well to the empirical site-frequency-spectrum (Figure S13). Both the 2-Epoch model and the 3-Epoch model resulted in a similar likelihood and fit well to the empirical site-frequency-spectrum (Figure S14, Table S15). However, the 2-Epoch model is a more reasonable model, as the parameters inferred from the 3-Epoch model are inconsistent with what we know about human populations. More specifically, an ancestral population size of >5 million is unreasonable, and the inferred bottleneck timing would be ~9 million years ago—well before human-chimp divergence.

Relevant scripts:

- D1_emit_all.sh
- See also: https://github.com/SexChrLab/Kenya_Selection_and_Demography/tree/master/Population_History

***Demographic simulations with SLiM 3***

We used a recently developed supervised machine learning method (*48*) to understand the history of selection at the *STC1* locus (Figure S15). In particular, we were interested in understanding: 1) whether the sweep involved a single adaptive allele (hard sweep) or multiple adaptive alleles (soft sweep), 2) if the sweep was soft, whether the alleles originated from recurrent new mutations (RNM) or from standing genetic variation (SGV), 3) the strength of selection (s), and 4) the timing of the onset of selection. Our method relies on a training set of summary statistics calculated from genomic data simulated under different evolutionary scenarios in SLiM 3 (*49*); the same summary statistics are then calculated for the test (focal) dataset. In particular, inferences are based on summary statistics describing patterns of nucleotide diversity, haplotype structure, and linkage disequilibrium, which are estimated across systematically varying genomic window sizes to capture sweeps across a wide range of selection strengths. For this analysis we focused on the 108 phased, high-coverage Turkana genomes and a ~1Mb region centered on the putatively selected region near *STC1* (chr8:23,350,029 - 24,424,864 in hg19 coordinates, which contains 3,538 biallelic polymorphic SNPs).

Using SLiM 3, we simulated genomic datasets under all three modes of selection: 1) hard sweeps, where an adaptive allele was introduced after the onset of selection; 2) RNM soft sweeps, where an adaptive allele was introduced after the onset of selection and a rate of mutation $\mu\alpha$ toward new adaptive alleles was set for the selected site; and 3) SGV soft sweeps, where a neutral allele was introduced and followed until it reached frequency $f_0$, at which point all copies of it in a population were converted into adaptive alleles. All simulations were run with uniform mutation and recombination rates of $\mu = r = 1.08 \times 10^{-9}$. RNM sweeps were simulated with an adaptive mutation rate of $8.3\times10^{-8} < \mu_\alpha < 4.2\times10^{-5}$ and SGV sweeps were simulated with an allele frequency at the onset of selection of $0.00004 < f_0 < 0.1$. $\mu_\alpha$ and $f_0$ were chosen randomly from a log-uniform distribution across their range of values. In all cases, we modeled a SNP under selection at the center of the simulated 1Mb region, and we assumed that adaptive alleles had selection coefficients > 0. Individuals homozygous for the wildtype allele were simulated to have fitness=1, individuals heterozygous for the adaptive allele had fitness=1 + hs, and individuals homozygous for the adaptive allele had fitness=1 + s. Selective sweeps were simulated with selection coefficients of $0.001 < s < 1.0$ under both codominant (h=0.5) and dominant (h=1.0) genetic architectures. In all simulations, when the sweep reached a frequency of 80%, we sampled 216 haplotypes from the population and calculated summary statistics from these data.

Simulations used a recapitation strategy for computational efficiency. First, forward-time dynamics of positive selection were simulated in SLiM 3 with an effective population size of 30,000 starting from a blank slate, without regard to the population's past. Afterward, the coalescence simulator msprime (*50*) was used to complete the ancestral genealogy of each simulation from the output of SLiM. The 2-Epoch demographic model was implemented during this coalescence step of simulation: the population size changed in the coalescence model from 15,000 to 30,000 at 7,000 generations prior to sampling. Lastly, we dropped neutral mutations on the resulting tree to generate the full dataset of neutral polymorphism around the sweep locus.

***Applying a convolutional neural network approach***

As input for the supervised machine learning model (a convolutional neural network, CNN), summary statistics of genetic variation were calculated for each simulation across base-pair subwindows for varying subwindow sizes. This analysis used 13 subwindow sizes and 13 center positions, with10 kb being the smallest subwindow size and the largest subwindow size covering the entire region. For each subwindow, seven summary statistics were calculated: the average nucleotide heterozygosity $\pi$, Tajima's D (*51*), the number of SNPs in the window, the number of distinct haplotypes in the window, $H_1$, $H_{12}$, and $H_2/H_1$ (*52*). Statistics were normalized linearly within pre-defined bounds; values outside the bounds were truncated to the minimum or maximum value for each statistic as appropriate. The same process of calculating and normalizing summary statistics was used to transform the empirical genotype data of SNPs around the *STC1* locus in order to provide it as input for the machine learning model.

For each of the three sweep modes (hard, RNM, and SGV sweeps), we used 5000 simulations as input for the CNN. A random 80% of the simulations were used for training and the rest were set aside for validation. We applied CNNs with two hidden convolutional layers as described in (*48*). One network was trained for inference of selection strength and another was trained for inference of sweep mode (hard, RNM, or SGV, respectively). 10 replicates of each network were trained on the same data for 50 epochs, using a batch size of 64. After training, models were used to make inferences on the empirical dataset. This process was repeated separately for simulations of codominant and dominant sweeps. CNNs were implemented in PyTorch and trained using the fastai v2.0 framework (*53*) with a one-cycle learning policy (*54*).

Finally, we compared our CNN results to an additional method that relies on different aspects of the sweep signature and assumes a different underlying evolutionary model. Specifically, we used CLUES, which performs a maximum likelihood estimation of the allele frequency trajectory and selection coefficient at a given site (*55*). To do so, it uses the posterior distribution of coalescence trees in the ancestral recombination graph (ARG) of the region. To run CLUES, we estimated the ARG with Relate, and we generated a sample of 3000 ARGs from their posterior distribution (*56*). CLUES was then run separately on the local coalescence tree for each of 30 candidate SNPs (focusing on the 30 loci closest to the center of the *STC1* candidate region). For each site, selection strength and allele frequency trajectory were calculated for the major allele (Figure S16). CLUES was then run using 1000 of the samples from Relate as a burn-in followed by thinning to 1 out of 100 of the remaining samples.

***Identifying traits under polygenic selection***

Given the polygenic nature of complex traits (*57*), we also searched for evidence of polygenic adaptation on cardiometabolic and renal system biomarkers using the approach described in (*43*). To do so, we used sets of SNPs previously associated with 29 urine and serum biomarkers via GWAS in UK

Biobank (Table S15). These GWAS summary statistics were derived specifically from meta-analyses conducted across diverse ancestries (downloaded from https://pan.ukbb.broadinstitute.org/).

To test for polygenic selection on a particular quantitative trait, we first obtained all 50kb windows associated with a given trait and summarized their genomic properties. We split the genome into non-overlapping 50 kb windows, and for each window calculated 1) the total number of SNPs, 2) the number of conserved SNP positions (i.e., those that fell within GERP conserved elements (*58*)), 3) the average recombination rate (using the 1000 Genomes Phase 3 genetic map), and 4) the median Fisher's combined score (FCS). The per-SNP FCS was calculated as the sum, over our three selection statistics, of $-\log_{10}$(rank of the statistic/number of SNPs tested). Because the PBS and XtX statistics were generally highly correlated, we also calculated the median FCS using just the |iHS| and XtX statistics so as not to overweight the two tests that focus on allele frequency differences.

We considered a 50 kb window to be associated with a given quantitative trait if the window contained a significant SNP ($p<5x10^{-8}$ from the meta-analysis across ancestries). After obtaining all 50 kb regions associated with a given trait, we generated a null distribution by randomly sampled x windows (x being the number of windows associated with the trait) among windows with a similar number of total SNPs, similar number of conserved SNPs, and similar recombination rate. To generate this null distribution, we only sampled from windows that were not associated with the focal trait, and we defined "similar" as within the same quartile of the total SNP, conserved SNP, or recombination rate distribution. We performed 1000 samples for a given trait, and then computed a p-value for each trait by calculating the proportion of resampled windows for which the median FCS value was higher than that observed for the real trait-associated windows. All p-values were adjusted for multiple testing and traits passing a 1% FDR were considered to be under polygenic selection (Table S15).

Relevant scripts:

- P1_UKBB_files.sh
- P2_polygenic_selection.R
- F_polygenic_selection.R

***Transcriptomic data generation***

Approximately 6mL of whole blood was collected in an EDTA tube by the THGP, and peripheral blood mononuclear cells (PBMCs) were then isolated using the SepMate protocol (STEMCELL Technologies). Isolated PBMCs were stored in RNALater and placed in a cooler box with frozen ice packs for 1-7 days, until the sample could be stored long-term at -20C. Samples were later thawed and extracted with the Zymo Quick-RNA kit. We also used ~10ul of whole blood from each EDTA tube to create a blood smear following standard protocols. This smear was fixed, subject to Wright-Giemsa staining (Camco brand stain pack), and viewed under a microscope; 100 total cells were counted to estimate the proportion of eosinophils, basophils, neutrophils, lymphocytes, and monocytes.

mRNA-seq libraries enriched for the 3' end of the mRNA molecules were prepared using TM3'seq (*59*). In brief, 10ul of input mRNA was used for first strand cDNA synthesis. This step was primed with the Tn5Me-B-30T oligo that binds to the polyA tail of mRNA molecules and has adapter-B sequence; together, these characteristics allow for 3' enriched libraries. Three rounds of PCR were used for cDNA amplification, followed by cDNA tagmentation using homemade Tn5 transposase pre-charged with adapter-A. Final library amplification was done for 12 PCR cycles using Illumina's i5 and i7 primer sequences that are complementary to the adapter-A and adapter-B sequences. As a result, only 3' cDNA fragments are amplified. The step by step TM3'seq protocol can be found in Suppl. File 1 of Pallares et. al. (*59*). Following the final library amplification step, 2µl of the final PCR amplification reactions were pooled by plate and size selected using a double-sided Agencourt AMPure XP bead (Beckman Coulter) cleanup approach. Pooled and cleaned libraries were visualized on an Agilent TapeStation, and sequenced using 100 bp SE reads on the Illumina NovaSeq S2 platform at the

Genomics Core Facility of the Lewis-Sigler Institute for Integrative Genomics at Princeton University. Protocols and subroutines for mRNA extraction, cDNA synthesis, and library preparation were implemented in the CyBio® FeliX liquid handling robot and are available upon request.

***Transcriptomic analyses***

Each mRNA-seq sample was trimmed for low quality bases and adapter contamination using cutadapt (*17*). Trimmed reads were then mapped to the human reference genome (hg38) using STAR (*60*) and filtered for unique mapping. Using our set of uniquely mapped reads, we counted the number of reads that overlapped each gene using HTSeq (*61*) and the GENCODE v25 GTF (https://www.gencodegenes.org/human/release_25.html). If a sample had fewer than 250,000 reads mapped to protein coding genes, we excluded it from further analyses.

We translated our raw count data into transcripts per million (TPM), and filtered the dataset to exclude non-protein coding genes as well as genes that were lowly expressed (median TPM<1). This filtering left us with 8438 genes. Next, we used the voom function in the R package limma (*62*) to normalize the count data. To remove variance attributed to technical batch effects, we used ComBat (*63*) to regress out sequencing lane. We then fit linear mixed effects models in the R package EMMREML to ask whether the expression of each gene was associated with lifestyle (urban or rural as defined in (*12*), and grouping pastoralists and non pastoralists together because of sample sizes and previous work (*12*)). All models included a pairwise genetic relatedness matrix derived from the array data and also controlled for fixed effects of age, the first PC of the genotype matrix, and cell type heterogeneity. More specifically, we created blood smears for all individuals included in the transcriptomic dataset and used a Wright-Giemsa stain approach to estimate the proportion of neutrophils, basophils, eosinophils, lymphocytes, and monocytes for each individual. We performed PCA on these 5-part differentials and included the first PC as a covariate in our differential expression analyses, because this PC explained 89.2% of the variance in cell type heterogeneity.

We extracted the p-value and effect size estimates for the lifestyle effect from each model and corrected for multiple hypothesis testing using a false discovery rate approach (*64*) and a 10% FDR cutoff (which left us with 490 differentially expressed (DE) genes). We then used gene set enrichment analyses (GSEA) (*65*) to ask whether certain biological pathways were overrepresented among the set of genes with the strongest evidence for differential expression. Specifically, we sorted our gene list by effect size (output from EMMREML) and used the "gseGO" function in the R package clusterProfiler (*66*) to perform GSEA.

Finally, we intersected our list of DE genes (n=490, FDR<10%) and a list of non-DE genes (n=7608, FDR>20%) with the set of SNPs tested for signatures of selection. We considered a SNP to be associated with a given gene if it fell within the gene body as well as within 100 kb of the transcription start or end site. We then ran linear models asking whether SNPs near DE versus non-DE genes differed in their iHS, PBS, or XtX statistics. All models controlled for gene expression level in the form of gene expression deciles (Table S19).

Relevant scripts:

- E1_process_RNAseq.R
- E2_DE_selection_overlap.R

***Ethics approval***

This study was approved by Princeton University's Institutional Review Board for Human Subjects Research (IRB #10237), Maseno University's Ethics Review Committee (MSU/DRPI/MUERC/00519/18), and Arizona State University's Institutional Review Board for Human Subjects Research (IRB ID: STUDY00004874). We also received county-level approval for research activities, and research permits from Kenya's National Commission for Science, Technology and Innovation (NACOSTI/P/18/46195/24671).

***Data and code availability***

Genomic data collected from indigenous groups is subject to their sovereignty. This includes their determination of uses that benefit the communities and/or align with their priorities, that protect against the risk of identification of subjects or groups, and that minimize uses with potentially stigmatizing interpretations. Therefore, we worked with study communities to establish a formal process for data sharing and secondary research proposals. Specifically, the data reported here are available to qualified researchers who sign a data use agreement approved by all stakeholders. This process involves: 1) submission of a proposal for secondary research that aligns with human subjects approvals and the general evolutionary and health-related scientific interests of the study communities, and that addresses data management and confidentiality as well as explicit benefits sharing (i.e., how access to genomic data and/or the proposed research will benefit study communities); 2) review of the proposal by a board comprised of representatives from Turkana communities, Kenyan scientists, and members of the THGP and ASU teams; and 3) individual and institutional approval of a data use agreement. The data are hosted on dbGap at accession numbers phs002219.v1.p1 and [TBD], and specific procedures for applying for data access can be found there.

The genomic data collected from the ASU team are hosted on dbGap under the accession number phs002219.v1.p1. This data is under general research use and requires a letter of collaboration.

Analysis code is available at https://github.com/AmandaJLea/Turkana_selection; https://github.com/SexChrLab/Kenya_Selection_and_Demography/tree/master/Population_History; and https://github.com/ayroles-lab/turkana-stc1.

### SUPPLEMENTARY FIGURES

**Figure S1. Map of current homelands/major areas for populations included in this study.** We note this map was created by the authors based on their personal knowledge as well as the literature, it is only meant to provide a general idea of group locations (rather than highly accurate and specific boundaries).

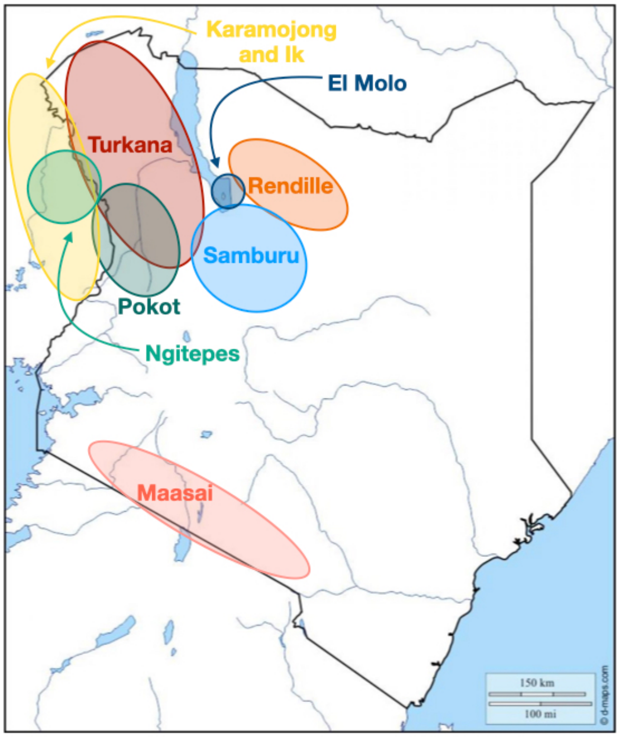

**Figure S2. Imputation accuracy in low/medium coverage whole genome sequencing (WGS) data is higher with a one panel approach.** (A-B) Scatterplots show the proportion of sites to impute for low/medium coverage samples versus the proportion of correctly imputed (concordant) genotypes estimated by cross-validation within IMPUTE2. The concordance estimates are derived from an imputation pipeline that uses (A) two reference panels (1000 Genomes and Turkana high coverage WGS data) or (B) one panel (Turkana high coverage WGS data). Blue dots represent samples from individuals of self-reported Turkana ancestry and yellow dots represent samples from individuals belonging to other groups. (C) Concordance estimates from the one panel versus two panel approach (dashed line represents x=y).

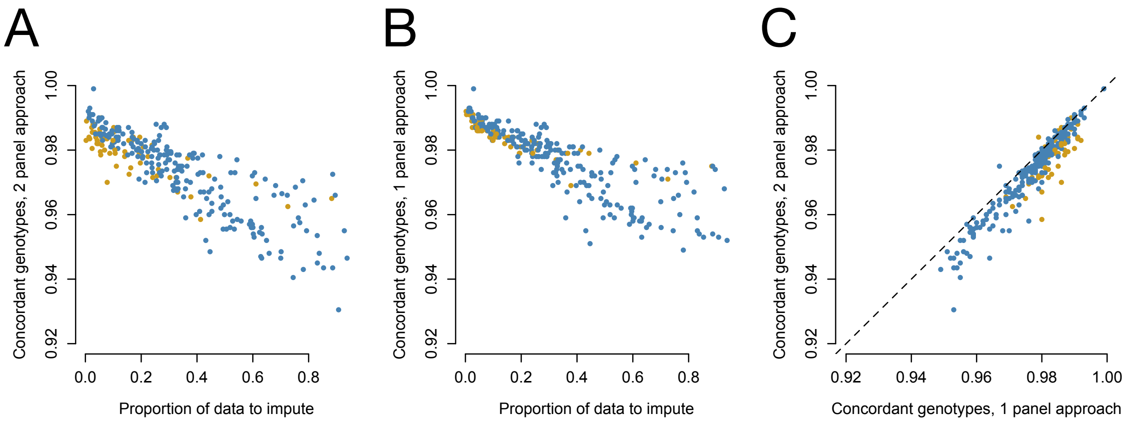

**Figure S3. Principal components analysis of whole genome sequencing data (focusing on samples included in this study).** (A) Loadings for the top two principal components (PCs), colored by self-reported ancestry group. Data are also divided by coverage group (high versus low/medium) and by sequencing location (Princeton versus ASU). Inset shows the percent variance explained (PVE) for each of the top 10 PCs. (B) Boxplots of principal components loadings broken down by self-reported ancestry group and by sequencing location. We do no observe any systematic biases in genetic variation as a result of sequencing location.

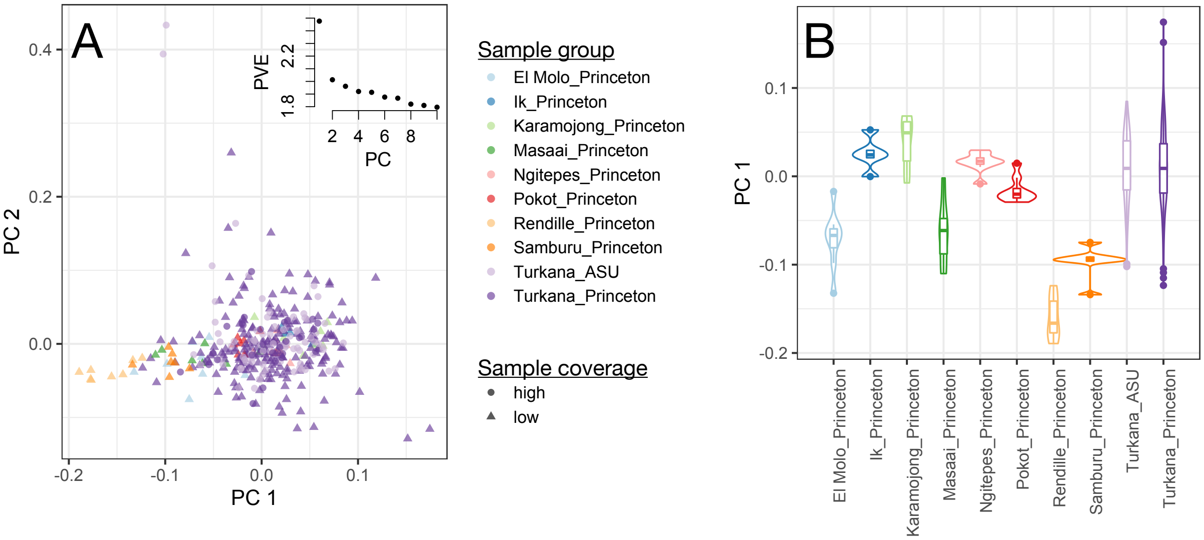

**Figure S4. Genetic variation does not systematically differ by sequencing location or coverage.** Y-axis shows the FDR-corrected p-value from a Wilcoxon signed-rank test. Each test asked whether there were differences between the principal component loadings from a PCA on the imputed, filtered genotype call set, focusing on contrasts between: 1) high coverage samples sequenced at Princeton versus ASU (blue dots) or 2) high versus low/medium coverage Turkana samples (yellow dots). Dashed line represents an FDR cutoff of 10%.

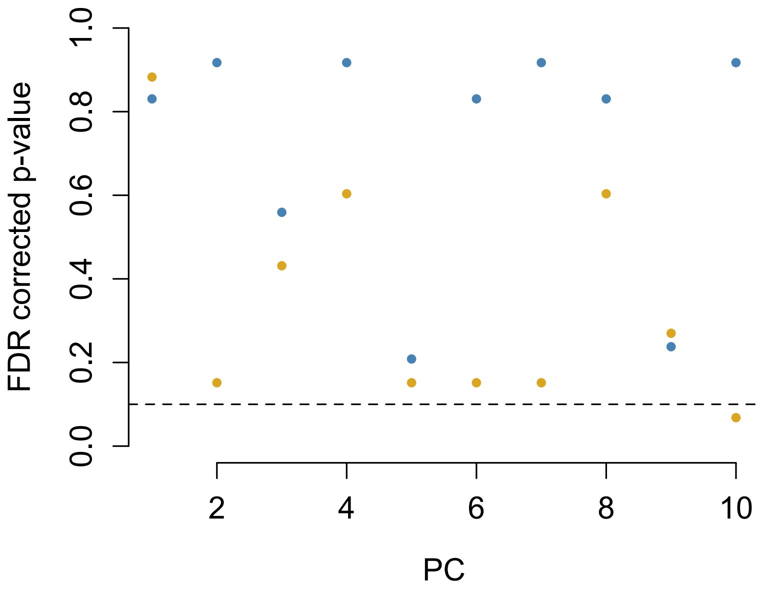

**Figure S5. Principal components analysis of array genotype data.** (A) Loadings for the top two principal components (PCs), colored by self-reported ancestry group (n=782). Inset shows the percent variance explained (PVE) for each of the top 10 PCs. (B-C) Boxplots of principal components loadings broken down by self-reported ancestry group.

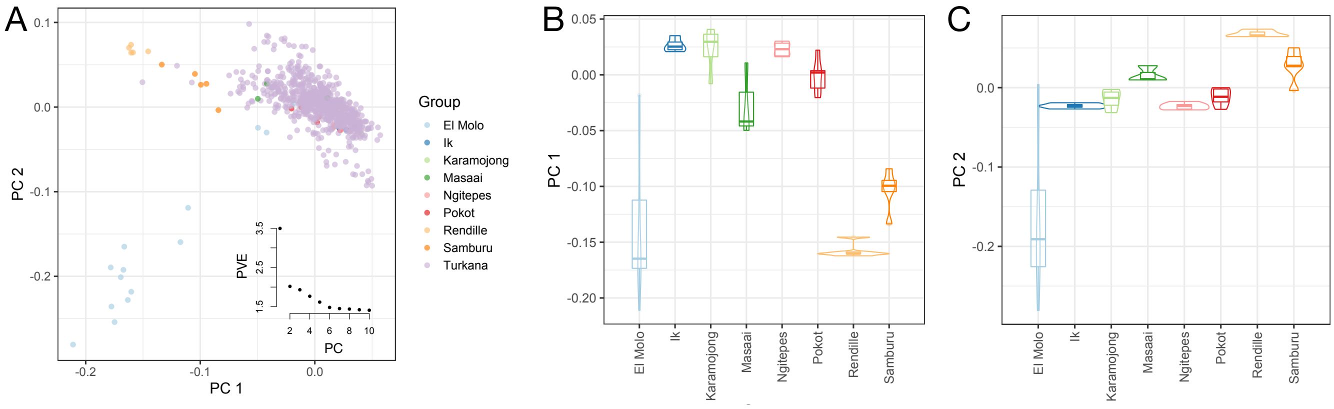

**Figure S6. Ancestry group and geography predict genetic variation.** Percent variance explained (PVE) by (A) self-reported ancestry group and (B) location for the top 10 principal components (PCs) of the array genotype data (n=782). PVE is equivalent to the $R^2$ from a linear model asking whether each PC was predicted by group or by longitude and latitude of the sampling location. Location analyses were conducted using only Turkana samples.

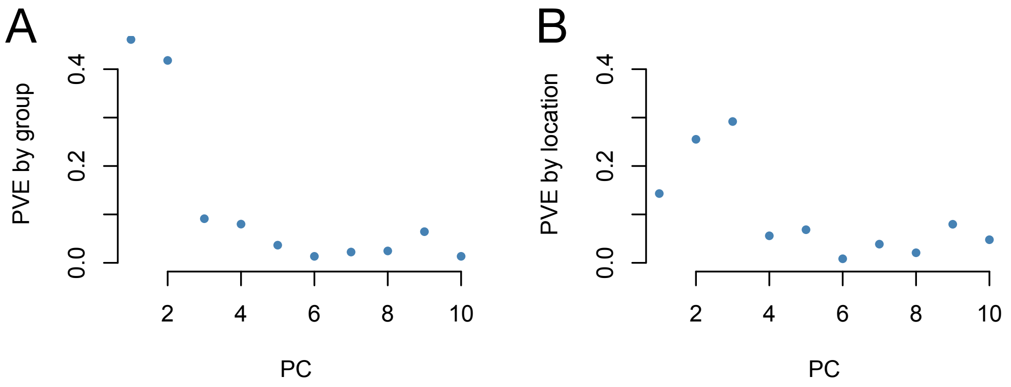

**Figure S7. Cross-validation error (CV) obtained from the program ADMIXTURE run with different values of K.** ADMIXTURE was run five times for each value of K with a different random seed.

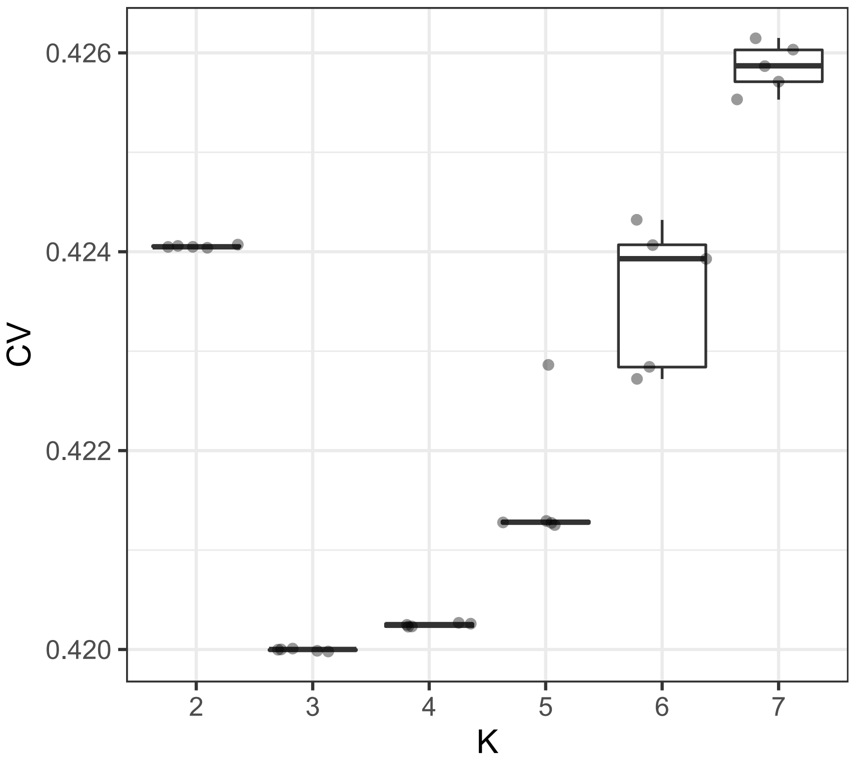

**Figure S8. Global ancestry analysis with ADMIXTURE.** Ancestry proportions inferred from the program ADMIXTURE run with different values of K. Specifically, panel A shows results from K=3 and panel B shows results from K=4 applied to the whole genome sequencing dataset. Panel C shows results for K=3 applied to the array dataset. Each bar represents an individual, and the height of the colored bar on the y-axis denotes the proportion of the genome assigned to a given ancestry component. Populations larger than n=10 were randomly downsampled for plotting. The number of colors correspond to the number of a prori defined ancestry components (K); colors are recycled across panels for visualization. Acronyms correspond to 1000 Genomes population codes as follows: CEU=Utah residents of Northern European ancestry, YRI=Yoruba; LWK=Luhya; MKK=Masaai.

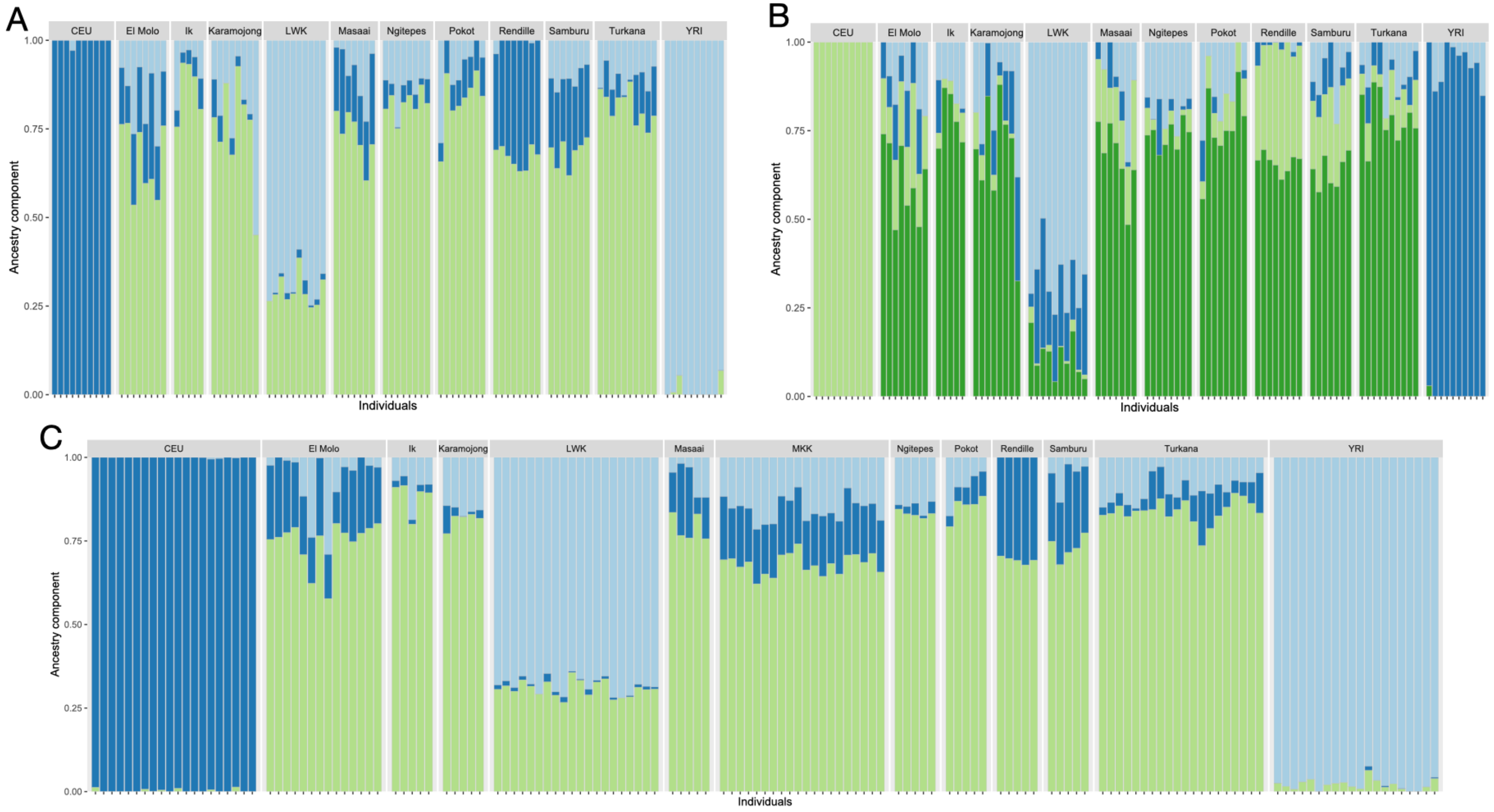

**Figure S9. Low levels of European ancestry in the Turkana.** A) Scatterplot shows the European ancestry proportions estimated from RFMix versus ADMIXTURE. This comparison focuses only on high coverage Turkana WGS samples, because RFMix requires phased genotype data. B) Distribution of European ancestry proportions estimated from ADMIXTURE (dashed line) versus RFMix (sold line).

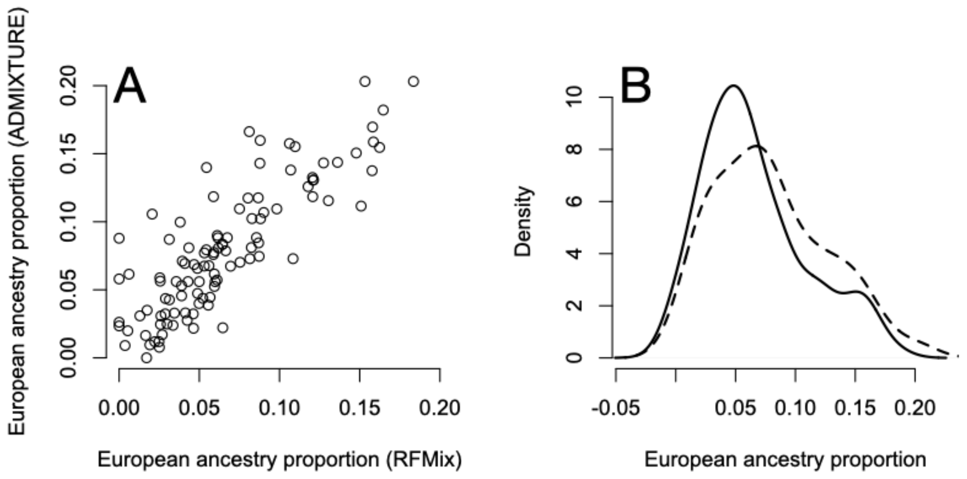

**Figure S10. Principal components analysis of whole genome sequencing data (combined with data from the 1000 Genomes Project).** (A) Loadings for the third and fourth principal components (PCs), colored by population. The loadings for the first two principal components are shown in the main text. (B-C) Boxplots of PC loadings broken down by population.

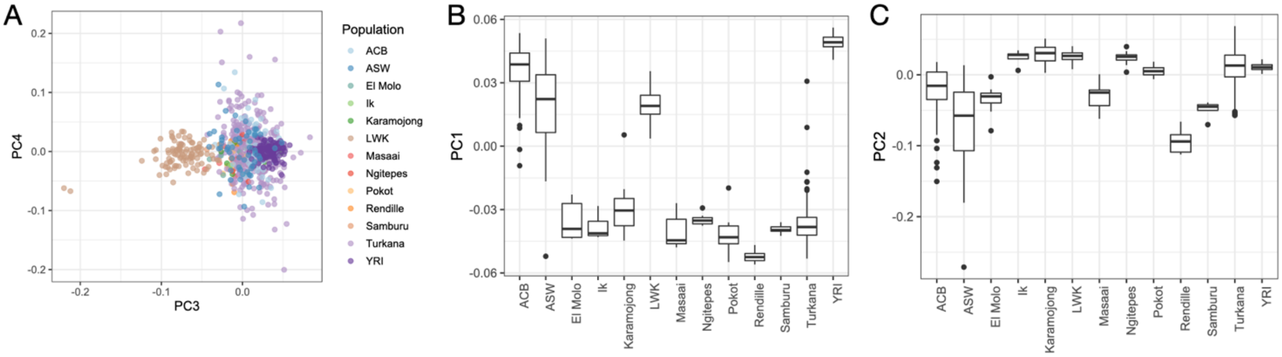

**Figure S11. HiC data from tissues that robustly express *STC1*.** Contact maps were obtained from Wang et al. 2018 (PMID: 30286773) and represent tissues that express *STC1* in the GTEx dataset. Coordinates are given in hg19 and the *STC1* gene body as well as the candidate region of interest are highlighted.

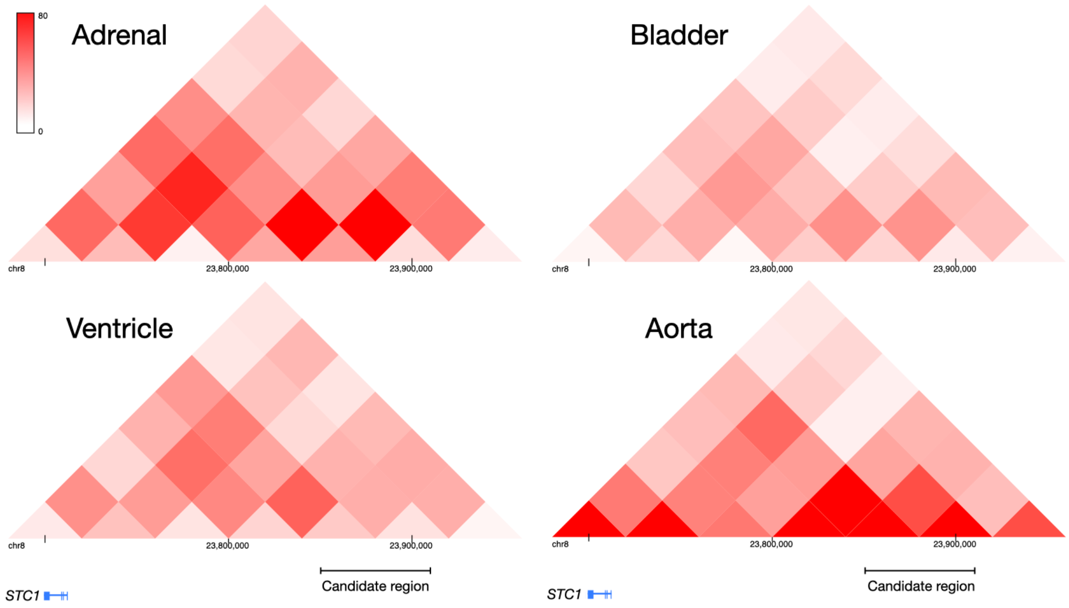

**Figure S12. Minimal genetic variation in the *STC1* candidate region in Europeans.** A) Minor allele frequencies for SNPs within the *STC1* candidate region for 1000 Genomes African (AFR) and European (EUR) populations. Coordinates are in hg19. Only 5 SNPs in the candidate region have an average MAF>5% in Europeans. Acronyms correspond to 1000 Genomes population codes as follows: ACB=African Caribbean; ASW=African American; CEU=Utah residents of Northern European ancestry; ESN=Esan; FIN=Finnish; GBR=British; IBS=Iberian; LWK=Luhya; MSL=Mende; TSI=Toscani; YRI=Yoruba. B) Allele frequencies for the same SNPs in the populations sampled as part of this study. The "minor" allele is matched to panel A, and therefore some SNPs have minor alleles >50% because they are the major allele in the focal East African populations.

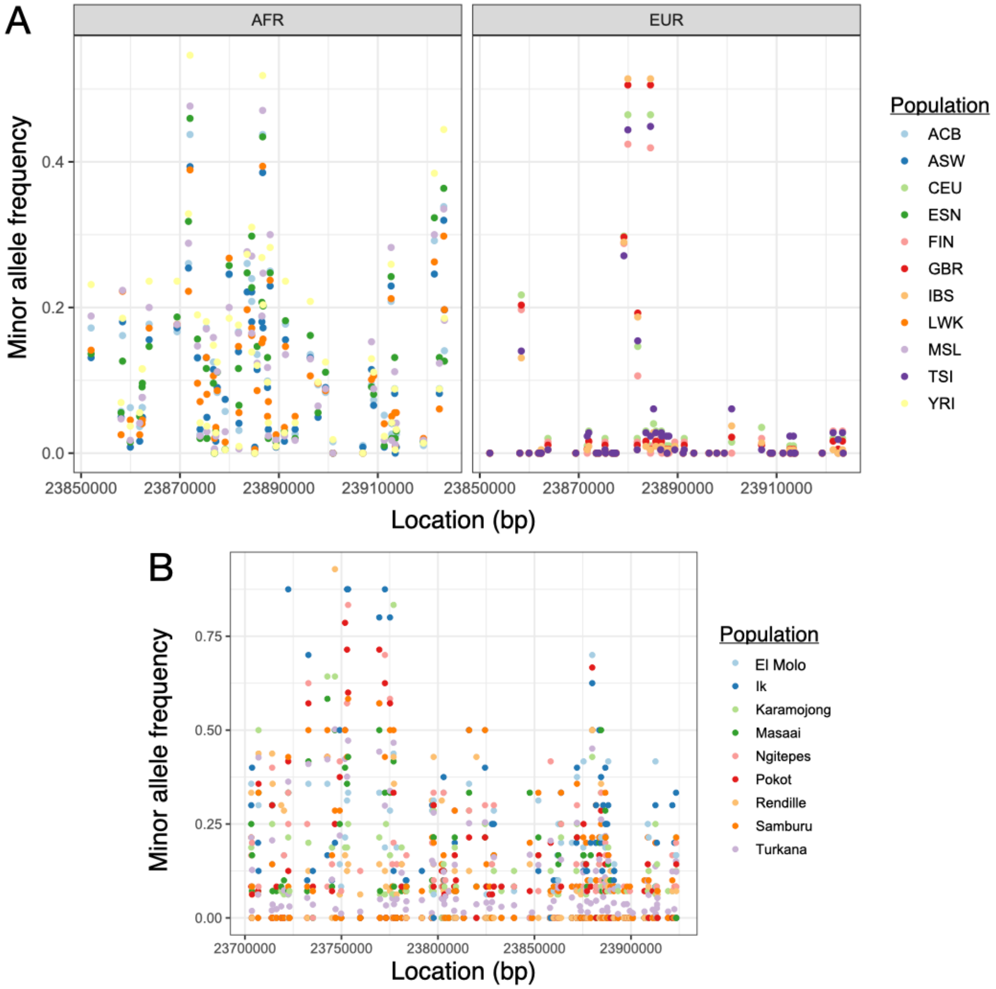

**Figure S13. Overview of demographic scenarios explored for the Turkana population.** Demographic inference was performed with DaDi and focused on SNPs within neutral regions that were genotyped in the high coverage Turkana sample set (n=108).

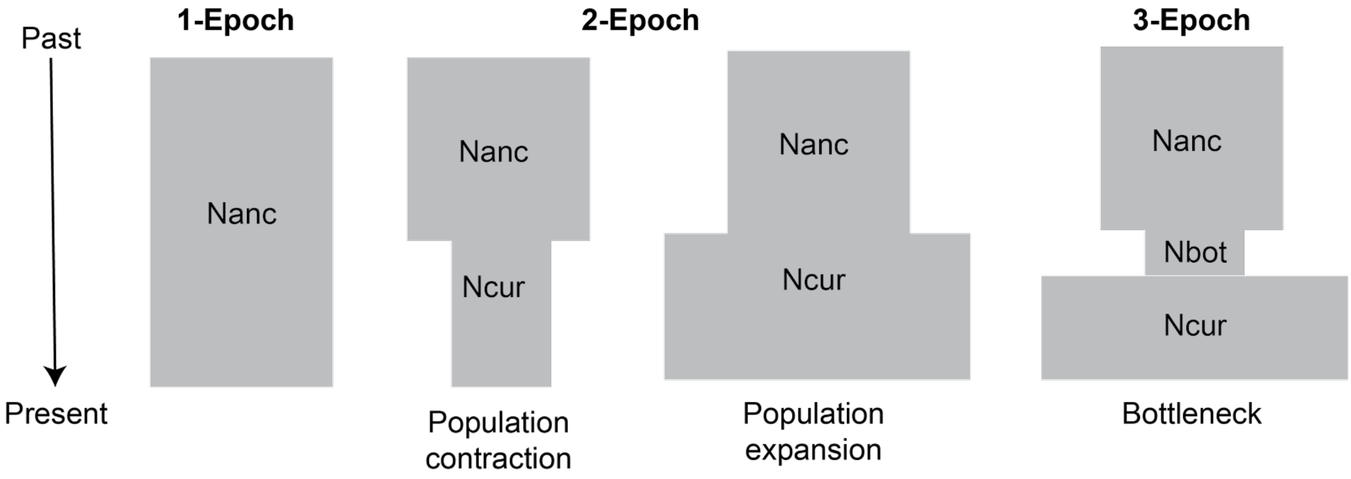

**Figure S14. Site frequency spectra** calculated from the empirical data (black) versus under different demographic scenarios (colors). The two and three epoch models provide a good visual fit to the empirical site frequency spectrum.

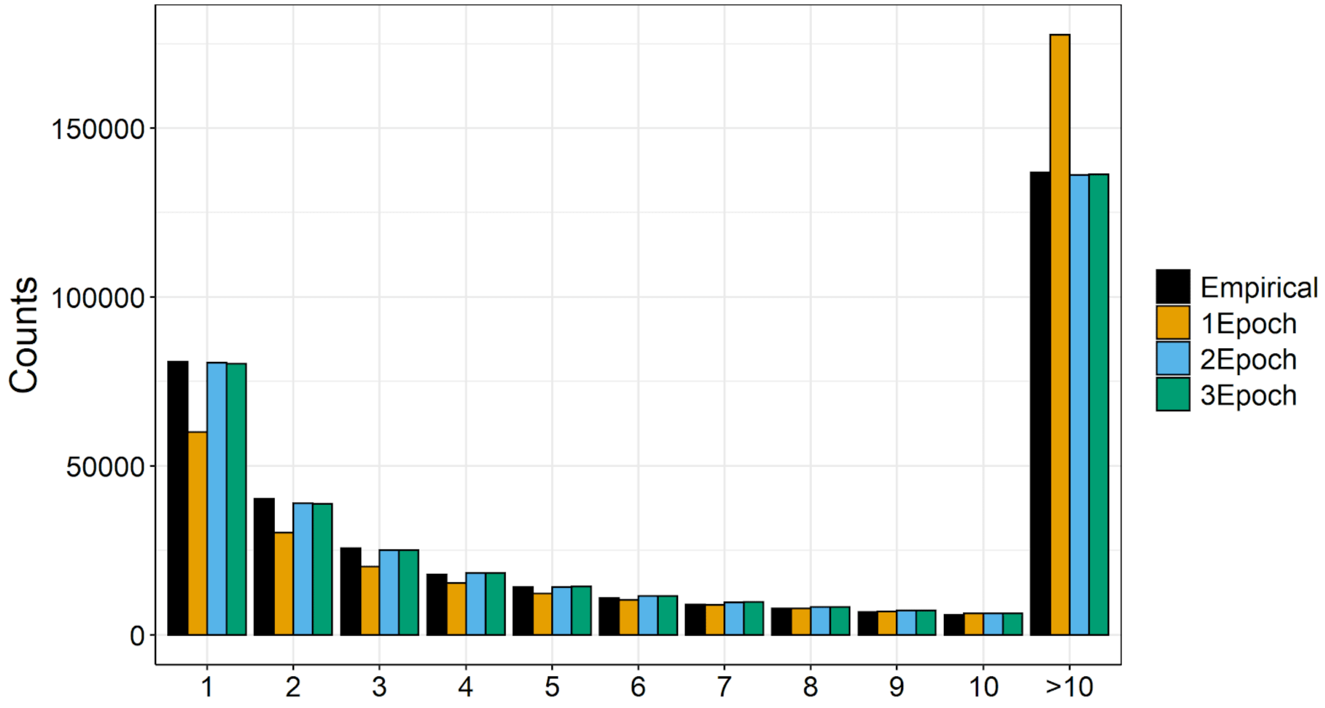

**Figure S15. Using a convolutional neural network (CNN) to infer the mode, strength, and timing of selection.** A) Diagram of machine learning-based method used to infer properties of the selective sweep at *STC1*. B) Results of applying 10 replicate CNNs trained to infer the selection coefficient (s) on the *STC1* locus, under the assumption that the selective sweep is either dominant or codominant.

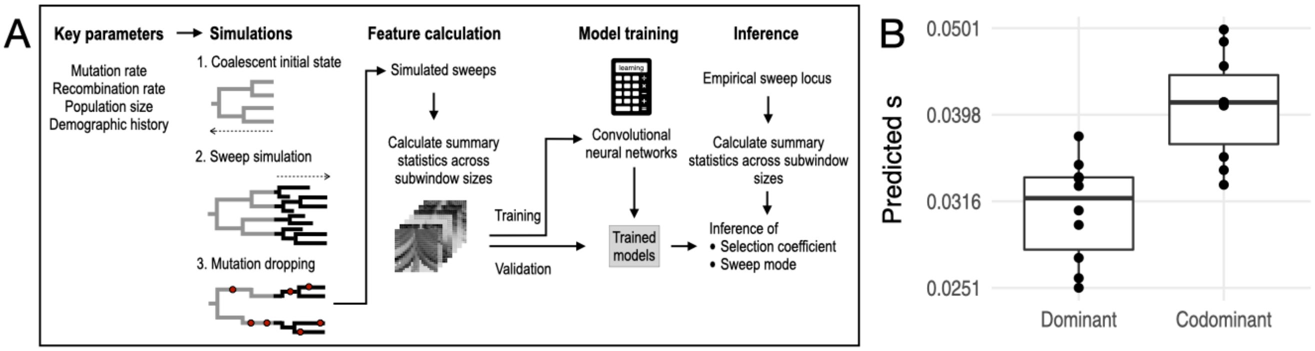

**Figure S16. Results of applying CLUES to SNPs within the *STC1* candidate region**. Brighter squares represent higher probability. The selection coefficient (s) estimated by CLUES is given in the plot.

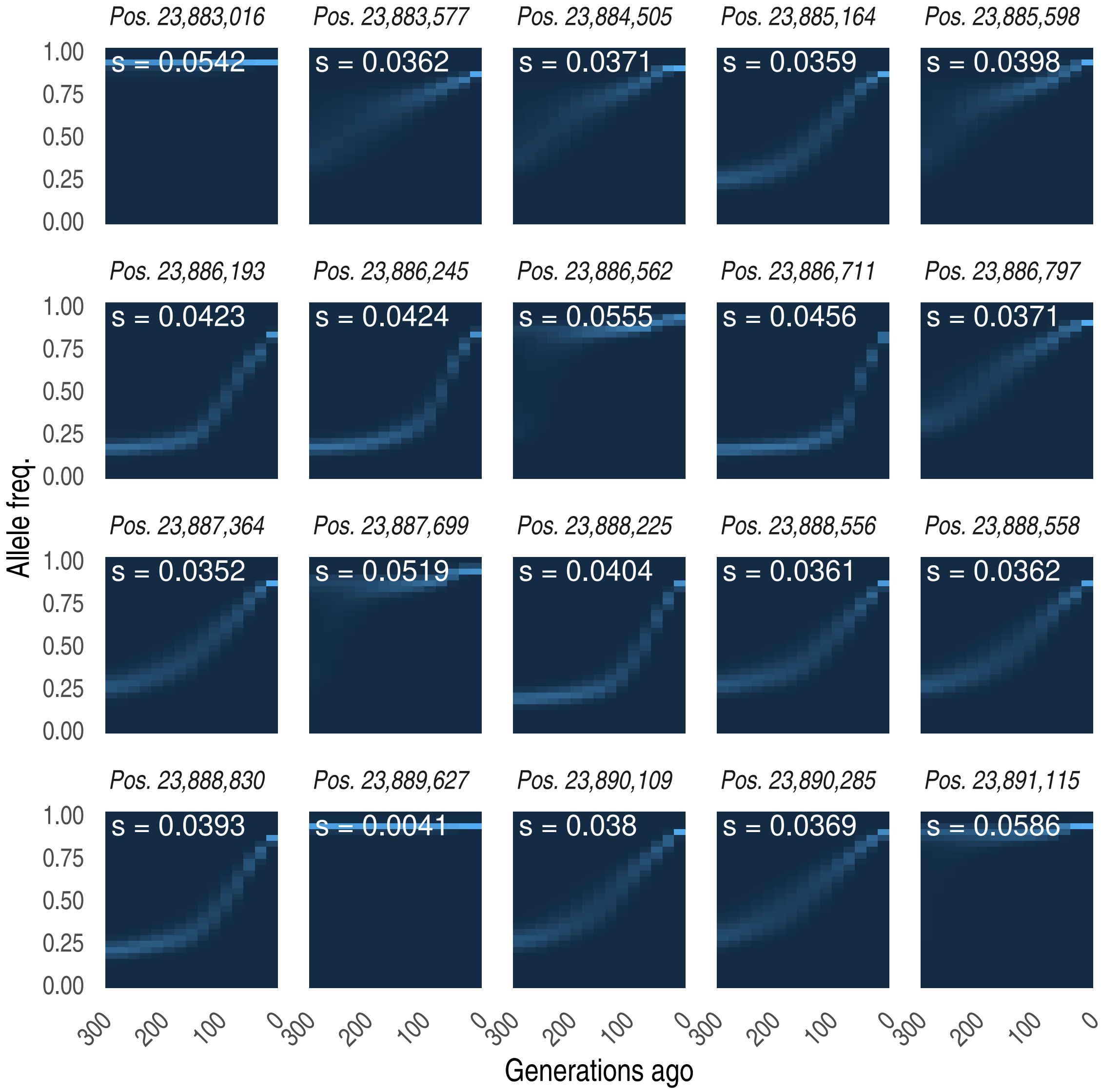

**Figure S17. Biological processes enriched within the set of genes differentially expressed by lifestyle** (urban versus rural Turkana; n=230). Gene set enrichment analysis was used to identify significant categories, and the default settings in the function "emapplot" from the R package enrichplot were used for plotting.

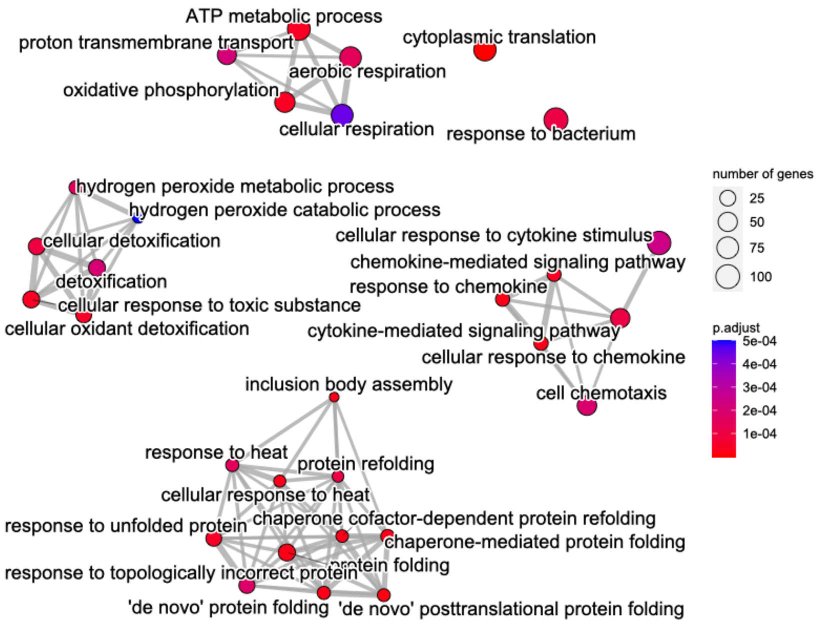

**Figure S18. Biological processes enriched within the set of genes differentially expressed by lifestyle** (urban versus rural Turkana; n=230). Gene set enrichment analysis was used to identify significant categories, and the default settings in the function “emapplot” from the R package enrichplot were used for plotting.

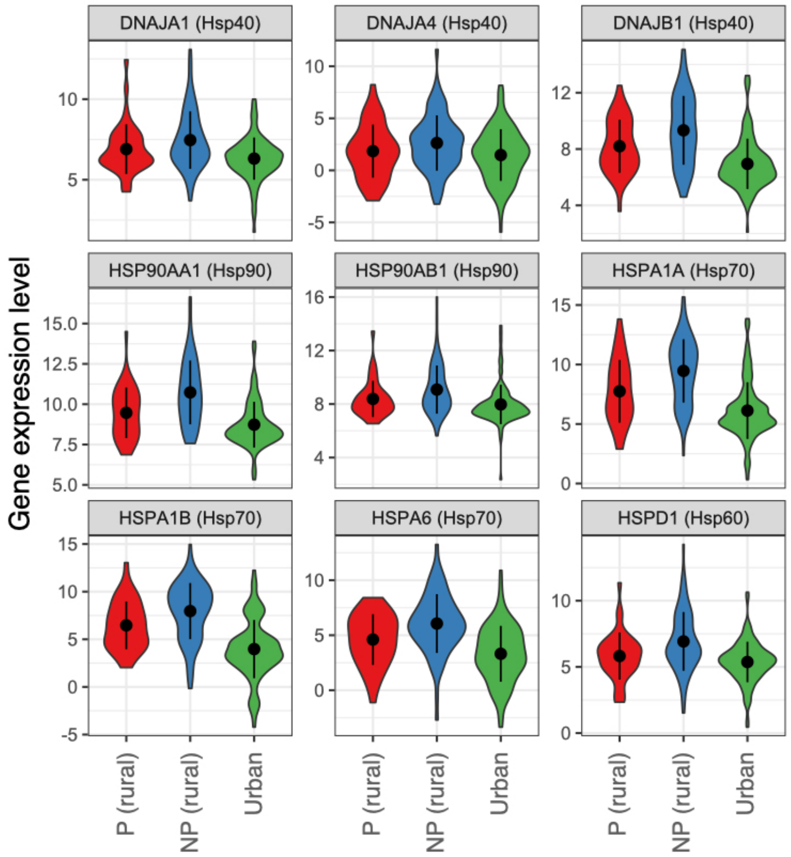

31. G. B. J. Busby, G. Band, Q. S. Le, M. Jallow, E. Bougama, V. D. Mangano, L. N. Amenga-Etego, A. Enimil, T. Apinjoh, C. M. Ndila, A. Manjurano, V. Nyirongo, O. Doumba, K. A. Rockett, D. P. Kwiatkowski, C. C. A. Spencer, Malaria Genomic Epidemiology Network, A. Vanderwal, A. Elzein, A. Nyika, A. Mendy, A. Miles, A. Diss, A. Kerasidou, A. Green, A. E. Jeffreys, B. MacInnis, C. Hughes, C. Moyes, C. Hubbart, C. Malangone, C. Potter, D. Mead, D. Barnwell, D. Jyothi, E. Drury, E. Somaskantharajah, E. Hilton, E. Leffler, G. Maslen, G. Busby, G. M. Clarke, I. Ragoussis, J. A. Garcia, J. Rogers, J. deVries, J. Shelton, J. Ragoussis, J. Stalker, J. Rodford, J. O'Brien, J. Evans, K. Rowlands, K. Cook, K. Fitzpatrick, K. Kivinen, K. Small, K. J. Johnson, L. Hart, M. Manske, M. McCreight, M. Stevens, M. Pirinen, M. Hennsman, M. Parker, M. SanJoaquin, N. Seplúveda, O. Cook, O. Miotto, P. Deloukas, R. Craik, R. Wrigley, R. Watson, R. Pearson, R. Hutton, S. Oyola, S. Auburn, S. Shah, S. Q. Le, S. Molloy, S. Bull, S. Campino, T. G. Clark, V. Ruano-Rubio, V. Cornelius, Y. Y. Teo, P. Corran, N. De Silva, P. Risley, A. Doyle, J. Evans, R. Horstmann, C. Plowe, P. Duffy, D. Carucci, M. Gottleib, A. Tall, A. B. Ly, A. Dolo, A. Sakuntabhai, O. Puijalon, A. Bah, A. Camara, A. Sadiq, A. A. Khan, A. Jobarteh, A. Mendy, A. Ebonyi, B. Danso, B. Taal, C. Casals-Pascual, D. J. Conway, E. Onykwelu, F. Sisay-Joof, G. Sirugo, H. Kanyi, H. Njie, H. Obu, H. Saine, I. Sambou, I. Abubakar, J. Njie, J. Fullah, J. Jaiteh, K. A. Bojang, K. Jammeh, K. Sabally-Ceesay, L. Manneh, L. Camara, L. Yamoah, M. Njie, M. Njie, M. Pinder, M. Jallow, M. Aiyegbo, M. Jasseh, M. L. Keita, M. Saidy-Khan, N. Ceesay, O. Rasheed, P. L. Ceesay, P. Esangbedo, R. Cole-Ceesay, R. Olaosebikan, S. Correa, S. Njie, S. Usen, Y. Dibba, A. Barry, A. Djimdé, A. H. Sall, A. Abathina, A. Niangaly, A. Dembele, B. Poudiougou, E. Diarra, K. Bamba, M. A. Thera, O. Doumbo, O. Toure, S. Konate, S. Sissoko, M. Diakite, A. T. Konate, D. Modiano, E. C. Bougouma, G. Bancone, I. N. Ouedraogo, J. Simpore, S. B. Sirima, M. Troye-Blomberg, A. R. Oduro, A. V. O. Hodgson, A. Ghansah, F. Nkrumah, F. Atuguba, K. A. Koram, M. D. Wilson, N. A. Ansah, N. Mensah, P. A. Ansah, T. Anyorigiya, V. Asoala, W. O. Rogers, A. O. Akoto, A. O. Ofori, D. Ansong, D. Sambian, E. Asafo-Agyei, J. Sylverken, S. Antwi, T. Agbenyega, A. E. Orimadegun, F. A. Amodu, O. Oni, O. O. Omotade, O. Amodu, S. Olaniyan, A.

Ndi, C. Yafi, E. A. Achidi, E. Mbunwe, J. Anchang-Kimbi, R. Mugri, R. Besingi, T. O. Apinjoh, V. Titanji, A. Elhassan, A. Hussein, H. Mohamed, I. Elhassan, M. Ibrahim, G. Kokwaro, T. Oluoch, A. Macharia, C. Newton, D. H. Opi, D. Kamuya, E. Bauni, K. Marsh, N. Peshu, S. Molyneux, S. Uyoga, T. N. Williams, V. Marsh, B. Nadjm, C. Maxwell, C. Drakeley, E. Riley, F. Mtei, G. Mtove, H. Wangai, H. Reyburn, S. Joseph, D. Ishengoma, M. Lemnge, T. Mutabingwa, J. Makani, S. Cox, A. Phiri, A. Munthali, D. Kachala, L. Njiragoma, M. E. Molyneux, M. Moore, N. Ntunthama, P. Pensulo, T. Taylor, R. Carter, D. Fernando, N. Karunaweera, R. Dewasurendra, P. Suriyaphol, P. Singhasivanon, C. P. Simmons, C. Q. Thai, D. X. Sinh, J. Farrar, L. Van Chuong, N. H. Phu, N. T. Hieu, N. T. H. Mai, N. T. N. Quyen, N. Day, S. J. Dunstan, S. E. O'Riordan, T. T. H. Chau, T. T. Hien, A. Allen, E. Lin, H. Karunajeewa, I. Mueller, J. Reeder, L. Manning, M. Laman, P. Michon, P. Siba, S. Allen, T. M. E. Davis, Admixture into and within sub-Saharan Africa (2016), doi:10.7554/eLife.15266.

32. P. Deelen, M. J. Bonder, K. J. van der Velde, H.-J. Westra, E. Winder, D. Hendriksen, L. Franke, M. A. Swertz, Genotype harmonizer: automatic strand alignment and format conversion for genotype data integration. *BMC Res. Notes*. **7**, 1–4 (2014).

33. B. K. Maples, S. Gravel, E. E. Kenny, C. D. Bustamante, RFMix: a discriminative modeling approach for rapid and robust local-ancestry inference. *Am. J. Hum. Genet.* **93**, 278–288 (2013).

34. B. F. Voight, S. Kudaravalli, X. Wen, J. K. Pritchard, A map of recent positive selection in the human genome. *PLoS Biol.* **4**, 0446–0458 (2006).

35. X. Yi, Y. Liang, E. Huerta-Sanchez, X. Jin, Z. X. P. Cuo, J. E. Pool, X. Xu, H. Jiang, N. Vinckenbosch, T. S. Korneliussen, H. Zheng, T. Liu, W. He, K. Li, R. Luo, X. Nie, H. Wu, M. Zhao, H. Cao, J. Zou, Y. Shan, S. Li, Q. Yang, Asan, P. Ni, G. Tian, J. Xu, X. Liu, T. Jiang, R. Wu, G. Zhou, M. Tang, J. Qin, T. Wang, S. Feng, G. Li, Huasang, J. Luosang, W. Wang, F. Chen, Y. Wang, X. Zheng, Z. Li, Z. Bianba, G. Yang, X. Wang, S. Tang, G. Gao, Y. Chen, Z. Luo, L. Gusang, Z. Cao, Q. Zhang, W. Ouyang, X. Ren, H. Liang, H. Zheng, Y. Huang, J. Li, L. Bolund, K. Kristiansen, Y. Li, Y. Zhang, X. Zhang, R. Li, S. Li, H. Yang, R. Nielsen, J. Wang, J. Wang, Sequencing of 50 human exomes reveals adaptation to high altitude. *Science*. **329**, 75–78 (2010).

41. J. Russell, M. Mascher, I. K. Dawson, S. Kyriakidis, C. Calixto, F. Freund, M. Bayer, I. Milne, T. Marshall-Griffiths, S. Heinen, A. Hofstad, R. Sharma, A. Himmelbach, M. Knauft, M. van Zonneveld, J. W. S. Brown, K. Schmid, B. Kilian, G. J. Muehlbauer, N. Stein, R. Waugh, Exome

sequencing of geographically diverse barley landraces and wild relatives gives insights into environmental adaptation. *Nat. Genet.* **48**, 1024–1030 (2016).
